# Antagonists targeting eEF2 kinase rescue multiple aspects of pathophysiology in Alzheimer’s disease model mice

**DOI:** 10.1101/2021.07.15.452522

**Authors:** Nicole P Kasica, Xueyan Zhou, Xin Wang, Wenzhong Yang, Helena R Zimmermann, Caroline E Holland, Elizabeth Koscielniak, Hanzhi Wu, Anderson O Cox, Jingyun Lee, Alexey G Ryazanov, Cristina M. Furdui, Tao Ma

**Affiliations:** Department of Internal Medicine, Gerontology and Geriatric Medicine, Wake Forest School of Medicine, Winston-Salem, North Carolina, USA; Department of Internal Medicine-Section on Molecular Medicine, Wake Forest University School of Medicine, Winston-Salem, NC 27157, USA; Comprehensive Cancer Center, Wake Forest Baptist Medical Center, Winston-Salem, NC 27157, USA; Department of Pharmacology, Rutgers Robert Wood Johnson Medical School, Piscataway, New Jersey, USA; Department of Physiology and Pharmacology, Wake Forest School of Medicine, Winston-Salem, North Carolina, USA; Department of Neurobiology and Anatomy, Wake Forest School of Medicine, Winston-Salem, North Carolina, USA

## Abstract

It is imperative to develop novel therapeutic strategies for Alzheimer’s disease (AD) and related dementia syndromes based on solid mechanistic studies. Maintenance of memory and synaptic plasticity relies on *de novo* protein synthesis, which is partially regulated by phosphorylation of eukaryotic elongation factor 2 (eEF2) via its kinase eEF2K. Abnormally increased eEF2 phosphorylation and impaired mRNA translation have been linked to AD. We recently reported that prenatal genetic suppression of eEF2K is able to prevent aging-related cognitive deficits in AD model mice, suggesting the therapeutic potential of targeting eEF2K/eEF2 signaling in AD. Here, we tested two structurally distinct small-molecule eEF2K inhibitors in two different lines of AD model mice after onset of cognitive impairments. Our data revealed that treatment with eEF2K inhibitors improved AD-associated synaptic plasticity impairments and cognitive dysfunction, without altering brain amyloid β (Aβ) and tau pathology. Furthermore, eEF2K inhibition alleviated AD-associated defects in dendritic spine morphology, postsynaptic density formation, protein synthesis, and dendritic polyribosome assembly. Our results may offer critical therapeutic implications for AD, and the proof-of-principle study indicates translational implication of inhibiting eEF2K for AD and related dementia syndromes.

**One Sentence Summary:** Treatment with eEF2K inhibitors and genetic eEF2K knockout improved cognitive deficits in Alzheimer’s disease model mice.

## Introduction

Alzheimer’s disease (AD) is an aging-related multifactorial neuronal disease characterized by dementia syndrome and distinct neuropathology including beta-amyloid (Aβ) deposits and tau tangles (*1*). Currently, there is no effective intervention for curing AD or slowing disease progression, and completed clinical trials to date have not succeeded in identifying a viable disease-modifying strategy (*2*). Investigation of molecular signaling pathways that go awry in AD may shed light on novel therapeutic strategies and diagnostic biomarkers for this devastating disease. A substantial body of evidence demonstrates that long-lasting forms of memory and synaptic plasticity require *de novo* protein synthesis (mRNA translation) (*3, 4*). Previous studies have shown impaired translational capacity and ribosomal function in the brains of human AD patients (*3, 5–7*), and recent works indicate that protein synthesis impairments may contribute to cognitive deficits in neurodegenerative diseases including AD, prion disease, and frontotemporal dementia (*8–10*).

Protein synthesis takes place in 3 phases: initiation, elongation, and termination. Numerous studies have focused on the initiation process, as it is usually considered the rate-limiting step during protein synthesis (*11*). Meanwhile, more than 95% of the energy and amino acids consumed during mRNA translation are used in the elongation phase (*11, 12*). Consistently, the elongation process is subject to tight control and modulation during cellular responses to nutrient and energy deficiency, which has been linked to multiple neurodegenerative diseases (*5, 13, 14*). Further, low cellular mRNA translational capacity in neuronal dendrites would require upregulation of both initiation and elongation processes to meet the need of new protein synthesis associated with maintenance of memory and synaptic plasticity (*15*). One critical mechanism regulating elongation involves eukaryotic elongation factor 2 (eEF2) which mediates the translocation step of elongation, catalyzing movement of tRNA from the ribosomal A-site to the P-site via GTP hydrolysis (*16*). Phosphorylation of eEF2 at Thr56 by its only known kinase eEF2K, disrupts peptide chain growth and consequently represses general protein synthesis (*16, 17*). Previous work has demonstrated abnormal hyper-phosphorylation of eEF2 in *post mortem* brain tissue from AD patients and in the brains of AD mouse models (*10, 18*). Further, we recently showed that reduction of eEF2K/eEF2 signaling through prenatal genetic manipulation improved translational capacity and prevented the development of aging-related cognitive deficits in AD model mice (*19*). Based on these findings, we hypothesized that pharmacological inhibition of eEF2K/eEF2 signaling is of therapeutic benefit for AD. Here, we tested the hypothesis by applying two structurally distinct small-molecule antagonists of eEF2K after onset of cognitive deficits in two lines of AD model mice. We took a multidisciplinary approach to determine the effects of the eEF2K inhibitors on multiple aspects of AD pathophysiology. Our findings may have critical therapeutic implications for AD and other aging-related dementia syndromes.

## Results

### eEF2K inhibitor A-484954 rescues cognitive deficits in AD model mice

To investigate potential therapeutic effects of eEF2K inhibition on AD-associated cognitive deficits, we treated aged APP/PS1 AD model mice (referred to as APP mice hereafter) and littermate wild type (WT) mice with A-484954 (referred to hereafter as AG), a selective small-molecule eEF2K antagonist (*20*). All studies here were performed in aged mice when cognitive deficits can be reliably observed in AD models (please refer to the method section for details on sources and age of the mice). Following a treatment dose previously reported (*21*), AG was applied for two weeks by using subcutaneous pellets containing the inhibitor or vehicle (Fig. 1A). The mice were then subjected to a series of behavioral tasks for assessment of cognitive function. We first performed the open field (OF) task to assess baseline anxiety-like behavior and general locomotor activity. We observed no significant differences among the four treatment groups in OF including ratio time spent in peripheral area of the arena, as well as travel distance and velocity (Fig. 1B, Fig. S1A and B). Next, we performed the novel object recognition (NOR) task to evaluate long-term recognition memory (*22*). WT mice treated with either AG or vehicle exhibited a preference for the novel object over the familiar object (Fig. 1C). In contrast, the APP mice treated with vehicle spent more time with the familiar object than the novel object, indicating a cognitive deficit and behavioral inflexibility (Fig. 1C). Notably, the APP mice treated with AG (APP + AG) showed performance comparable to that of WT mice, spending significantly more time with the novel than with the familiar objects (Fig. 1C). Additionally, AG treatment did not alter weight of the mice (Fig. S1C).

**Fig. 1.**
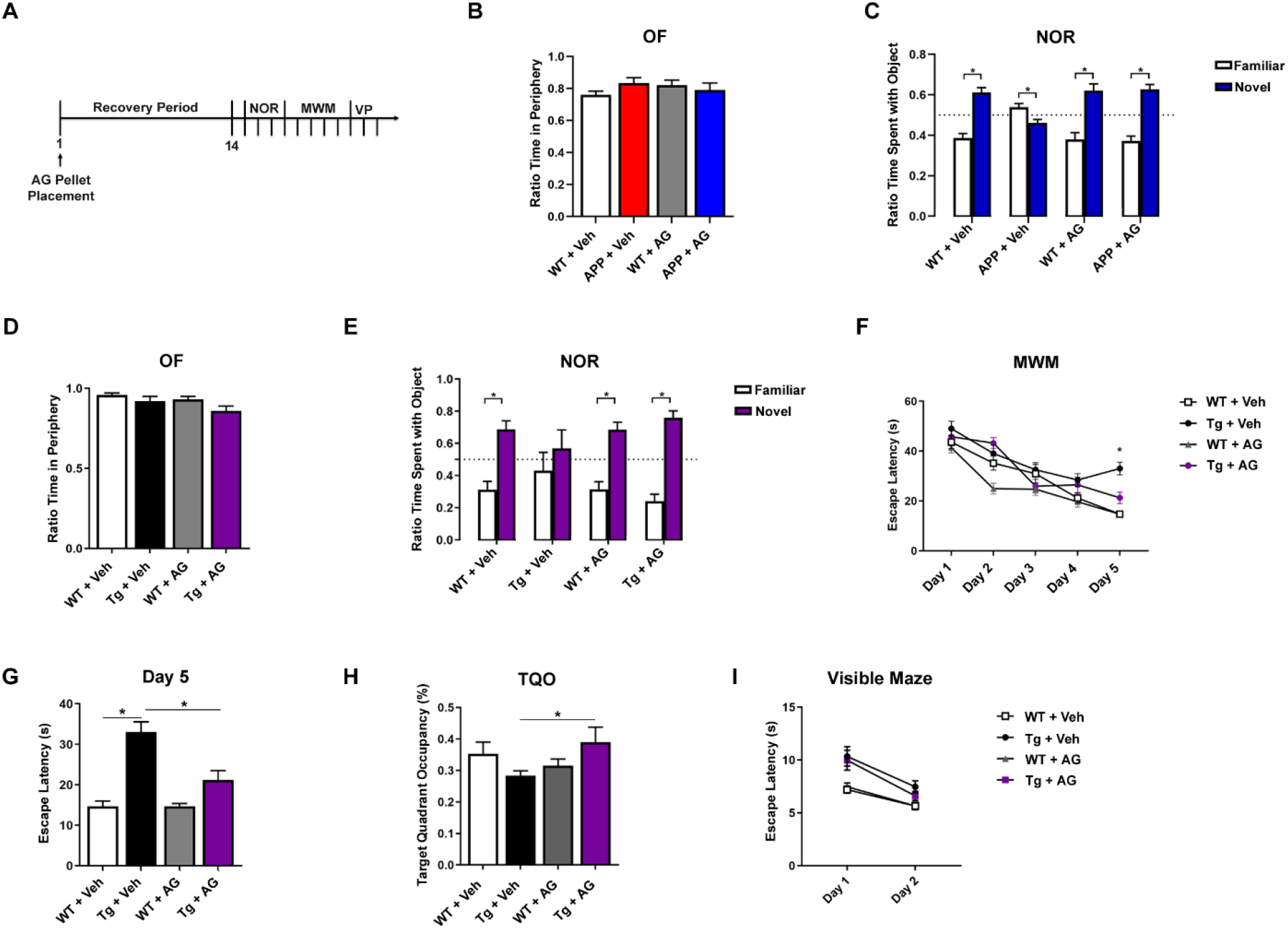
Treatment with eEF2K inhibitor A-484954 (AG) rescues cognitive deficits in APP/PS1 and Tg19959 AD model mice. **(A)** Treatment schematic showing placement of pellet containing AG or vehicle and subsequent behavioral tasks. **(B)** Ratio of time spent in the periphery for the OF test. (WT + Veh, *n* = 12; APP + Veh, *n* = 6; WT + AG, *n* = 8; APP + AG, *n* = 7). **(C)** Percentage of time spent with familiar (white) and novel (blue) objects in the NOR task during the testing phase. Preference of less than 50% indicates cognitive impairment. (WT + Veh, *n* = 12; APP + Veh, *n* = 6; WT + AG, *n* = 8; APP + AG, *n* = 7). Statistical preference for novel or familiar object: WT + Veh, \**p* < 0.0001; APP + Veh, *p* = 0.0107; WT + AG, \**p* = 0.0002; APP + AG, \**p* < 0.0001, unpaired *t* test. **(D)** Ratio of time spent in the periphery for the OF test (WT + Veh, *n* = 11; Tg + Veh, *n* = 10; WT + AG, *n* = 13; Tg + AG, *n* = 10). **(E)** Percentage of time spent with familiar (white) and novel (purple) objects in the NOR task during the testing phase. Preference of less than 50% indicates cognitive impairment. (WT + Veh, *n* = 6; Tg + Veh, *n* = 5; WT + AG, *n* = 6; Tg + AG, *n* = 7). Statistical preference for novel or familiar object: WT + Veh, \**p* = 0.0005; Tg + Veh, *p* = 0.4140; WT + AG, \**p* = 0.0002; Tg + AG, \**p* < 0.0001, unpaired *t* test. **(F)** Escape latency (s) over 5 days of training in the hidden platform MWM. Four trials/day, 5 days. (WT + Veh, *n* = 13; Tg + Veh, *n* = 11; WT + AG, *n* = 16; Tg + AG, *n* = 12). WT + Veh versus Tg + Veh, \**p* < 0.0001; WT + Veh versus Tg + AG, \**p* = 0.0484; Tg + Veh versus Tg + AG, \**p* = 0.0002, 1-way ANOVA with Tukey’s post hoc test. *F* = 24.08. **(G)** Escape latency on day 5 of MWM training. WT + Veh versus Tg + Veh, \**p* < 0.0001; Tg + Veh versus Tg + AG, \**p* = 0.0002, 1-way ANOVA with Tukey’s *post hoc* test. *F* = 24.08 **(H)** Percentage of time spent in target quadrant during probe trial phase of MWM task. (WT + Veh, *n* = 13; Tg + Veh, *n* = 11; WT + AG, *n* = 16; Tg + AG, *n* = 12). Tg + Veh versus Tg + AG, \**p* =0.05, unpaired *t* test. **(I)** Escape latency (s) over 2 days in the visible maze task. (WT + Veh, *n* = 13; Tg + Veh, *n* = 11; WT + AG, *n* = 16; Tg + AG, *n* = 12). Error bars represent + SEM.

We further examined whether the rescuing effects of eEF2K inhibition described above can be replicated in a different rodent model of AD. We thus applied the AG compound or vehicle to the Tg19959 (referred to as Tg hereafter) AD model mice (*23*). We did not observe any differences among the four treatment groups in the OF task (Fig. 1D, Fig. S1D and E). Consistent with the results from the experiments with the APP mice, impairments of recognition memory displayed in Tg mice (assessed by NOR test) were significantly improved with AG treatment (Fig. 1E). It is worth mentioning that Tg mice treated with vehicle, unlike vehicle-treated APP mice, did not spend more time with familiar objects, which might be due to the aging effects (see method section). We further used the hidden-platform Morris water maze (MWM) task to assess the performance on spatial learning and memory (*22*). Compared to WT mice treated with vehicle or AG, Tg mice treated with vehicle exhibited longer escape latency times, indicative of a cognitive impairment (Fig. 1F and G). Importantly, impairments of spatial learning and memory associated with Tg mice were alleviated by treatment with AG, as indicated by decreased escape latency and improved target quadrant occupancy (TQO) (comparable to Tg mice treated with vehicle) in the probe trial (Fig. 1F-H). Distance traveled and velocity of movement during the probe trial of MWM were not altered across all groups (Fig. S1F and G). Additionally, we examined potential memory-independent effects of AG (e.g. swimming ability, vision and motivation) through the visible maze task. No differences in latency to locate the visible platform were observed across all groups (Fig. 1I). We also did not observe alterations on weight and gross morphology of the hippocampus among treatment groups (Fig. S1H and I). Biochemical experiments demonstrated that increased levels of hippocampal eEF2 phosphorylation in AD model mice were blunted with AG treatment (Fig. S1J-L). Additionally, brain tissue was subjected to high performance liquid chromatography (HPLC) for measurement of AG concentration. AG-treated mice demonstrated the presence of AG in the brain, suggesting that the compound is able to cross the blood-brain barrier (Fig. S1M). Taken together, these results show that treatment with small-molecule eEF2K inhibitor AG rescues AD-associated learning and memory deficits.

### Aβ deposition and tau phosphorylation are not affected by the A-484954

We next investigated potential mechanisms through which pharmacological inhibition of eEF2K might confer protective effects on the AD model mice. We examined Aβ pathology with an immunohistochemical approach and found similar levels of Aβ plaque deposition in hippocampus of Tg mice treated with vehicle or AG (Fig. 2A-C). We also utilized ELISA to measure level of brain Aβ 1-40 or Aβ 1-42 and found no differences in Tg mice treated with AG or vehicle (Fig. 2D and E) or between APP mice treated with AG or vehicle (Fig. 2F and G). Furthermore, inhibition of eEF2K with AG did not affect levels of phosphorylated tau in either APP or Tg mice (Fig. S2A-F). Together, these results suggest that pharmacological eEF2K inhibition may improve cognitive function in AD model mice independently of regulation on Aβ or tau pathology.

**Fig. 2.**
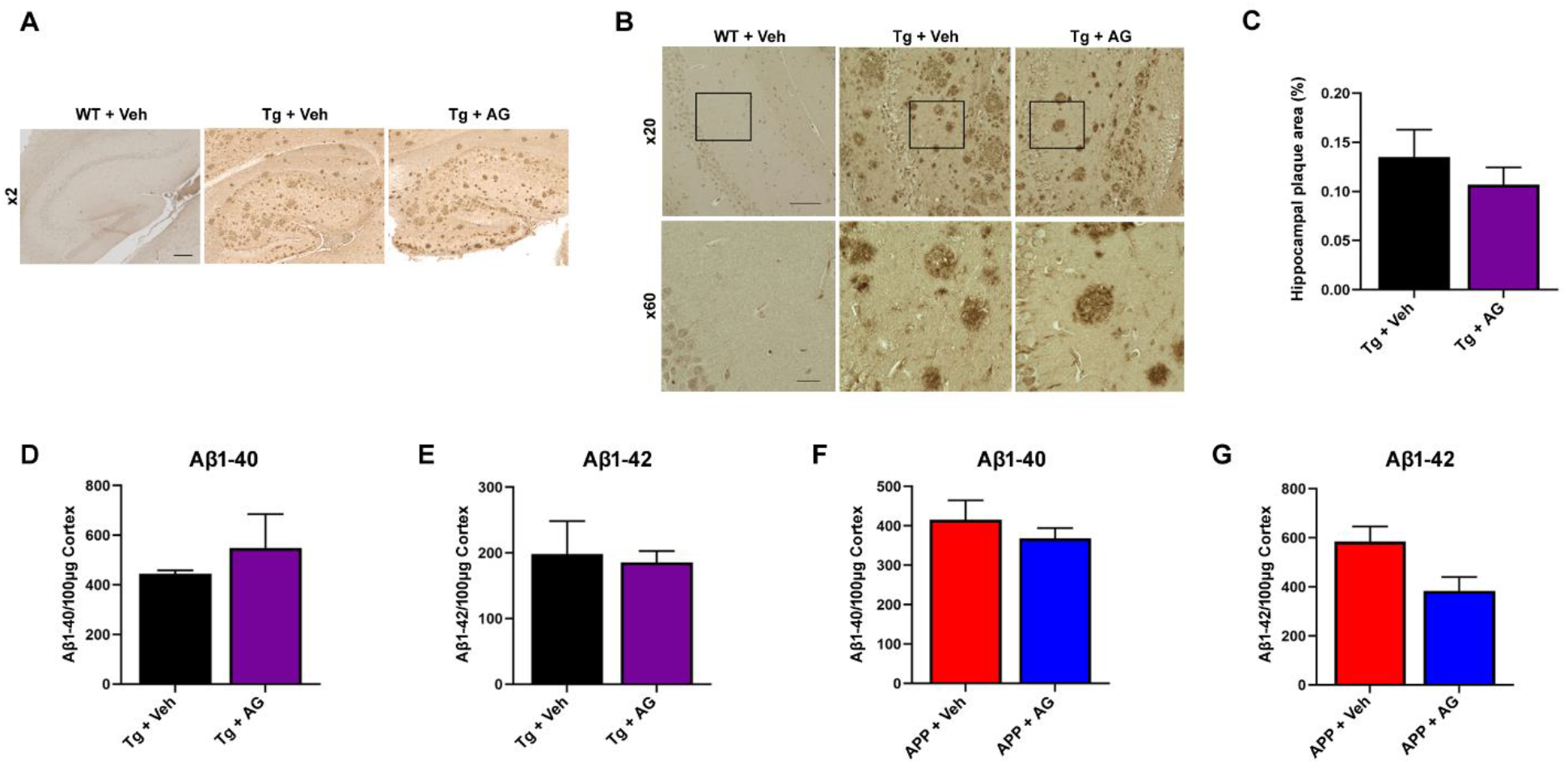
Amyloid plaque deposition is not altered with treatment of eEF2K inhibitor AG. **(A)** Representative images of hippocampal plaque deposition in WT + Veh, Tg19959 + Veh, and Tg19959 + AG mice at x2 magnification. Scale bar: 50 μm **(B)** Representative images of hippocampal plaque deposition in WT + Veh, Tg + Veh, and Tg + AG mice. Insets are shown at x60 magnification. Scale bars: 100 μm (x20); 20 μm (x60). **(C)** Percentage of hippocampal area covered in amyloid plaques in Tg+ Veh (*n* = 10 sections) and Tg + AG (*n* = 8). *p* = 0.45, unpaired *t* test. **(D)** ELISA showed no differences in levels of Aβ 1-40 or Aβ 1-42 **(E)** in Tg + Veh and Tg + AG forebrain tissue. *n* = 6. *p* = 0.55 for Aβ 1-40, *p* = 0.80 for Aβ 1-40, unpaired *t* test. **(F)** ELISA showed no differences in levels of Aβ 1-40 or Aβ 1-42 **(G)** in Tg + Veh and Tg+ AG forebrain tissue *n* = 6. *p* = 0.51 for Aβ 1-40, *p* = 0.06 for Aβ 1-42, unpaired *t* test. Error bars represent + SEM.

### A-484954 mitigates hippocampal protein synthesis deficits in AD model mice

Previous work has shown a marked decrease in translational capacity in the brains of AD patients and AD model mice (*19, 24–26*). We measured levels of *de novo* protein synthesis in mouse hippocampus using the surface sensing of translation (SUnSET), a non-radioactive puromycin end-labeling assay (*8, 19, 27*). Consistent with previous studies, hippocampal *de novo* protein synthesis (indicated by puromycin labeling) was significantly reduced in Tg mice compared with the WT group (Fig. 3A and B). Protein synthesis defects in the hippocampus of Tg mice were significantly improved with AG treatment, as indicated by improved puromycin labeling (Fig. 3A and B).

**Fig. 3.**
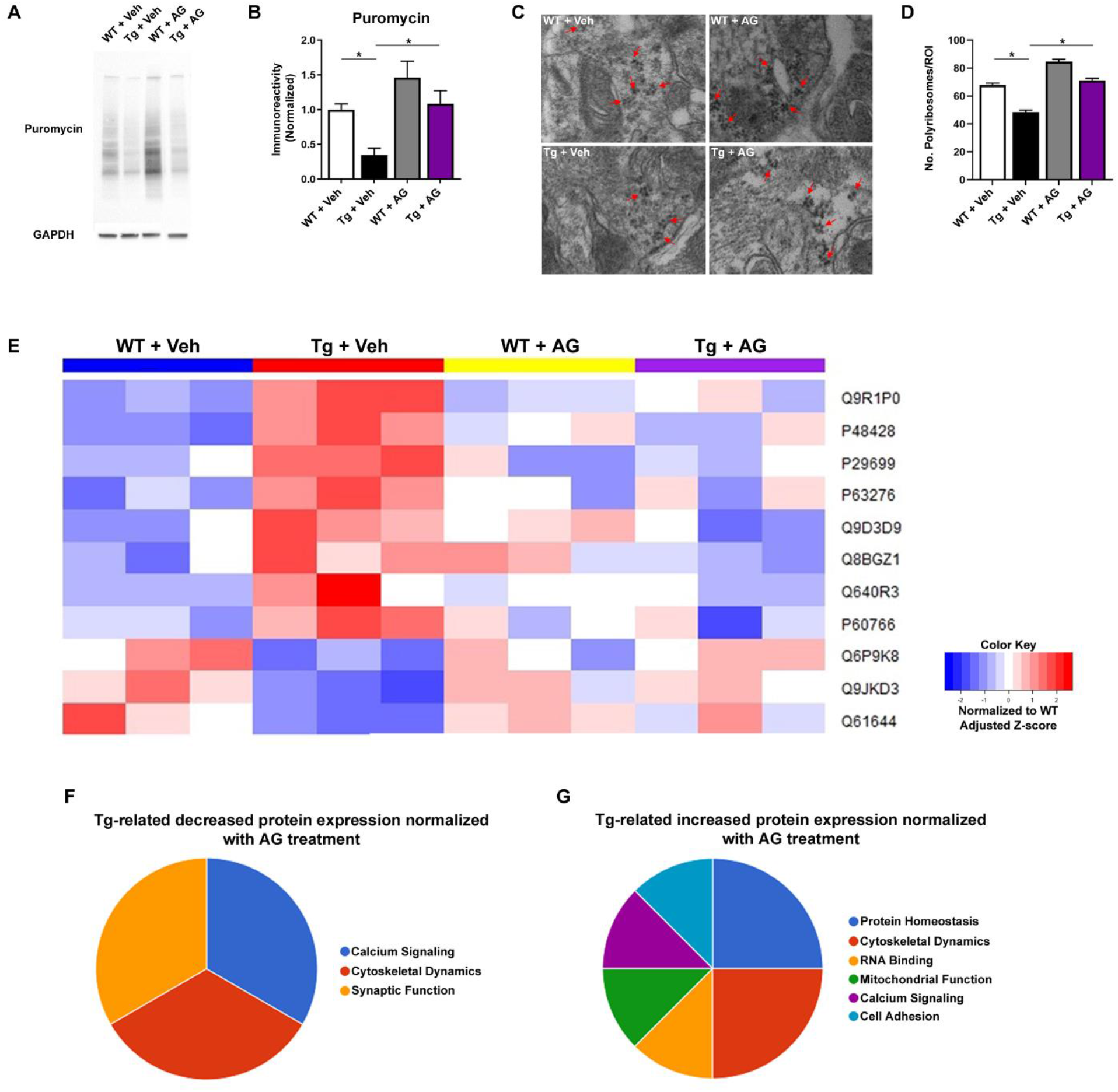
Treatment with AG mitigates protein synthesis deficits in hippocampus of an AD model mice. **(A)** Representative images from the SUnSET assay. Image shows 10-250 kDa range. **(B)** Quantification of *de novo* protein synthesis based on SUnSET assay. (WT + Veh, *n* = 6 mice; Tg + Veh, *n* = 6 mice; WT + AG, *n* = 6 mice; Tg + AG, *n* = 6 mice). WT + Veh versus Tg + Veh, \**p* < 0.05; Tg + Veh versus Tg + AG, \**p* = 0.02, 1-way ANOVA with Tukey’s post hoc test. *F* = 7.739. **(C)** Representative TEM images for CA1 polyribosomes. Arrows indicate polyribosomes. *n* = 3 mice per treatment group. Original magnification, x18,500. Scale bar = 500 nm. **(D)** Number of polyribosomes per ROI. WT + Veh versus Tg + Veh, \**p* < 0.0001; Tg + Veh versus Tg+ AG, \**p* < 0.0001, 1-way ANOVA with Tukey’s *post hoc* test. *F* = 99.63. Error bars represent + SEM. **(E)** Heat map showing patterns of protein expression in the brain of WT + Veh, Tg + Veh, WT + AG, and Tg + AG mice, reported as normalized deviation from the mean. **(F)** Functional classification of the decreased expression of proteins in AD mice that were restored with the eEF2K inhibitor treatment. *n* = 3 mice per genotype. **(G)** Functional classification of the increased expression of proteins in AD mice that were restored with eEF2K inhibitor treatment. *n* = 3 mice per genotype. More details are included in Table 1.

Polyribosomes are clusters of ribosomes engaged in active, ongoing translation; increased polyribosome count implies greater translational capacity and has been associated with the process of synaptic plasticity and memory formation (*28, 29*). We utilized transmission electron microscopy (TEM) to investigate whether inhibition of eEF2K activity via AG affected polyribosome assembly in area CA1 of the hippocampus. Compared with the WT group, Tg mice demonstrated a significant decrease in the number of polyribosomes, indicating impaired mRNA translation (Fig. 3C and D). Notably, Tg mice treated with AG exhibited increased number of hippocampal dendritic polyribosomes compared to the vehicle treatment group (Fig. 3C and D). These findings are consistent with the SUnSET data, suggesting improvement of *de novo* protein synthesis in AD mice with eEF2K inhibitor treatment.

**Table 1.** Top proteins dysregulated in Tg brains that are normalized with AG treatment.

| Ascension ID | Protein name | Function | Fold change (Tg v WT) | Fold change (Tg + AG v WT) |
| --- | --- | --- | --- | --- |
| Q9JKD3 | Secretory carrier-associated membrane protein 5 | Calcium Signaling | 0.76 | 1.24 |
| Q61644 | Protein kinase C and casein kinase substrate in neurons protein 1<sup>#</sup> | Cytoskeletal Dynamics | 0.82 | 1.92 |
| Q6P9K8 | Caskin-1<sup>#</sup> | Synaptic Function | 0.71 | 0.94 |
| Q9R1PO | Proteasome subunit alpha type-4 | Protein Homeostasis | 1.61 | 1.13 |
| P48428 | Tubulin-specific chaperone A | Cytoskeletal Dynamics | 3.49 | 1.13 |
| P29699 | Alpha-2-HS-glycoprotein | Protein Homeostasis | 1.98 | 0.95 |
| P63276 | 40S ribosomal protein S17 | RNA Binding | 1.39 | 0.94 |
| Q9D3D9 | ATP synthase subunit delta, mitochondrial | Mitochondrial Function | 2.48 | 0.93 |
| Q8BGZ1 | Hippocalcin-like protein 4 | Calcium Signaling | 1.20 | 0.99 |
| Q640R3 | Hepatocyte cell adhesion molecule | Cell Adhesion | 1.60 | 1.03 |
| P60766 | Cell division control protein 42 homolog | Cytoskeletal Dynamics | 1.27 | 0.98 |

<sup>#</sup>*p* < 0.08 in Tukey's post-hoc test between Tg + Veh and Tg + AG

We further examined potential effects of eEF2K inhibition on brain protein expression with unbiased mass spectrometry (MS)-based proteomic analysis. We were particularly interested in proteins whose expression is dysregulated (increased or decreased) in the brain of AD mice and restored with the eEF2K inhibitor treatment. Briefly, of all the proteins identified (1736 in total), 11 proteins in total met these criteria with statistical significance (Table 1, see method for details). A heat map was generated to summarize alterations of all these proteins (Fig. 3E). We further entered the protein information into the “Uniprot consortium” (https://www.uniprot.org) to uncover the functional classification of these 11 proteins (Fig. 3F and G). In brief, levels of 3 proteins were decreased in Tg mice (compared to WT) and improved by the eEF2K inhibitor treatment. Functionally, these proteins are involved in calcium signaling, cytoskeletal dynamics, and synaptic function (Fig. 3F and G, Table 1). On the other hand, there were 8 proteins whose expression levels increased in Tg (compared to WT) and were blunted with inhibition of eEF2K activity. By function, these proteins belong to 6 broad categories: protein homeostasis, cytoskeletal dynamics, RNA binding, mitochondrial function, calcium signaling, and cell adhesion (Fig. 3F and G, Table 1). We performed additional biochemical experiments to examine effects of AG treatment on other signaling pathways controlling protein synthesis (potentially linked to eEF2K/eEF2 signaling and AD) including AMP-activated protein kinase (AMPK), eukaryotic initiation factor 2α (eIF2α), and mammalian target of rapamycin (mTOR) (*14, 30*). We did not observe any significant effects by AG (Fig. S3). These findings indicate that treatment with the eEF2K inhibitor could rescue AD-associated deficiency in protein synthesis.

### A-484954 alleviates deficiency of spine morphology and synaptic structure in AD mice

Loss of synapses correlates robustly with memory impairments in human AD patients and AD animal models (*31–33*). Furthermore, *de novo* protein synthesis can affect spine density, morphology, and synapse strength (*4, 15*). Using the rapid Golgi-Cox staining protocol (*34*), we assessed spine density in apical dendrites of area CA1 of the hippocampus. First, we did not observe a significant difference in overall spine density across all four groups (Fig. 4A and B). We further analyzed differences in spine morphology according to published guidelines on classification of “mature” (branched, mushroom, and stubby) and “immature” (thin, long-thin, and filopodia) spines (*34*). Compared to the WT group, APP mice treated with vehicle showed a decrease of mature spines and an increase of immature spines (Fig. 4C and D). Markedly, the spine morphology deficits (for both mature and immature spines) were restored with the eEF2K inhibitor AG treatment (Fig. 4C and D). Furthermore, we applied TEM to assess postsynaptic densities (PSDs), which are located at the heads of spines and are critical for synaptic function (*35, 36*). Ultrastructural analysis of hippocampal area CA1 revealed decreased number of PSDs in Tg mice compared to the WT group (Fig. 4E and F). Notably, Tg mice treated with AG demonstrated a significant increase in number of PSDs compared to Tg vehicle-treated mice (Fig. 4E and F), suggesting a rescue effect of eEF2K inhibition. Additionally, we measured the length of the active zone of the PSDs, and found that there was a decrease in length of PSDs in Tg mice treated with vehicle compared to vehicle-treated WT mice (Fig. 4G). This deficit was alleviated in Tg mice treated with AG, suggesting a rescue effect of AG treatment (Fig. 4G). Taken together, these results suggest treatment with eEF2K inhibitor AG rescues defects of dendritic spine morphology, as well as PSD formation and length in AD model mice.

**Fig. 4.**
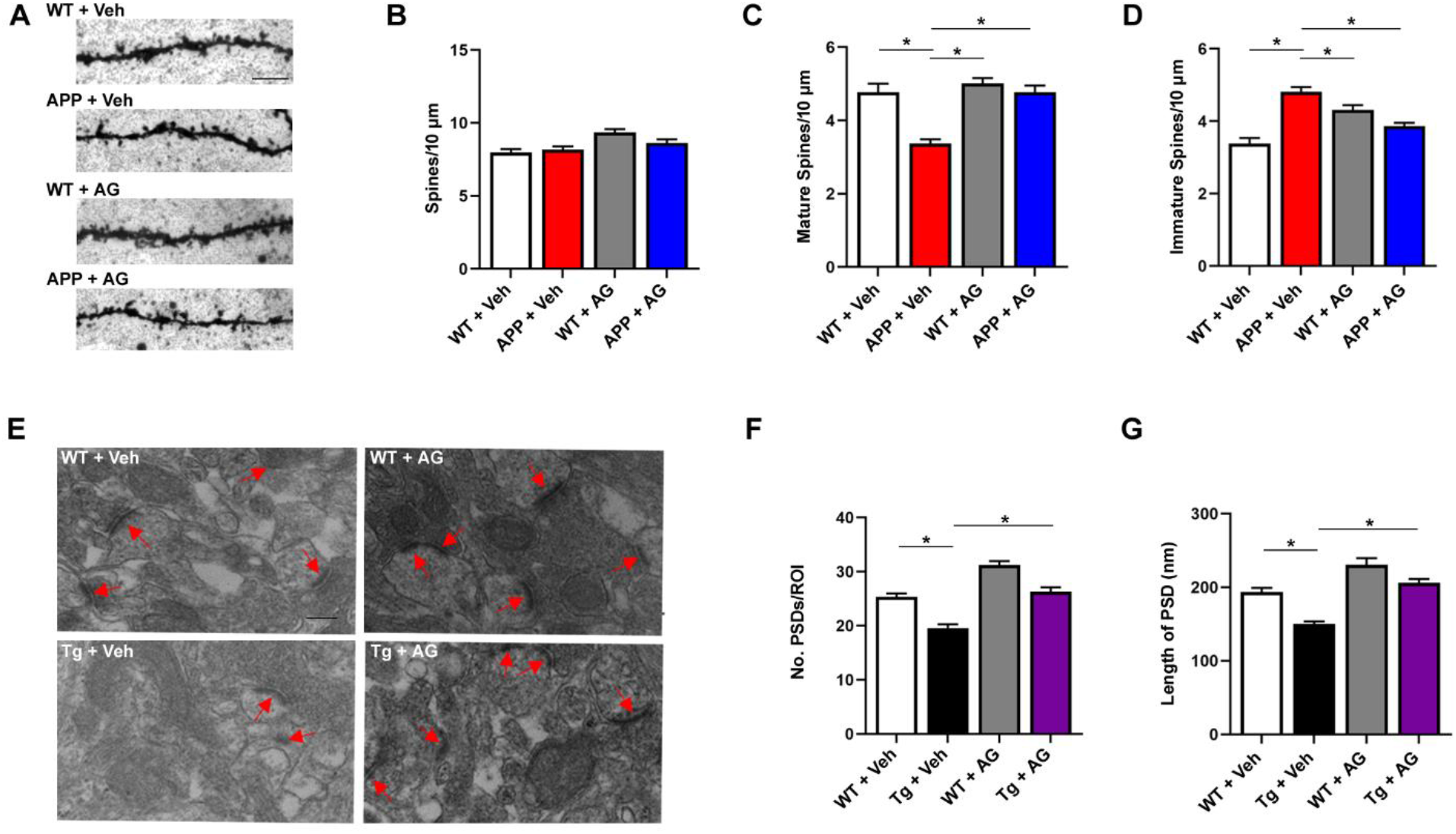
Treatment with eEF2K inhibitor AG alleviates defects in spine morphology and synaptic structure in an AD model mice. **(A)** Representative images from Golgi-Cox stain of area CA1 dendritic spines. Original magnification, x100. Scale bar: 12 μm. **(B)** Analysis of the total CA1 spine density per 10 μm. WT + Veh, *n* = 52 dendrites; APP + Veh, *n* = 73; WT + AG, *n* = 73; APP + AG, *n* = 55. **(C)** Measurement of mature spine density. Branched, mushroom, and stubby spines were classified as mature. APP + Veh mice had a significantly lower density of mature spines than WT + Veh, WT + AG, and APP + AG. APP + Veh versus WT + Veh, \**p* < 0.0001; APP + Veh versus WT + AG, \**p* < 0.0001; APP + Veh versus APP + AG, \**p* < 0.0001, 1-way ANOVA with Tukey’s *post hoc* test. *F* = 91.70. **(D)** Measurement of immature spine density. Filopodia, thin, and long-thin spines were classified as immature. APP + Veh mice had a significantly higher density of immature spines than WT + Veh, WT + AG, and APP + AG. APP + Veh versus WT + Veh, \**p* < 0.0001; APP + Veh versus WT + AG, \**p* < 0.0001; APP + Veh versus APP + AG, \**p* < 0.0001, 1-way ANOVA with Tukey’s post-hoc test. *F* = 91.70. **(E)** Representative TEM images for CA1 PSDs. *n* = 3 mice per genotype. Original magnification, x18,500. Scale bar: 500 nm. **(F)** Number of PSDs per ROI. WT + Veh versus Tg19959 + Veh, \**p* < 0.0001; Tg19959 + Veh versus Tg19959 + AG, \**p* < 0.0001, 1-way ANOVA with Tukey’s *post hoc* test. *F* = 43.31. **(G)** Length of active zone of PSD in nm. WT + Veh versus Tg + Veh, \**p* < 0.0001; Tg + Veh versus Tg + AG, \**p* < 0.0001, 1-way ANOVA with Tukey’s post hoc test. *F* = 30.34. Error bars represent + SEM.

### eEF2K inhibitor NH125 alleviates cognitive dysfunction and hippocampal LTP impairments in APP/PS1 AD model mice

Hippocampal long-term potentiation (LTP) is a major form of synaptic plasticity and cellular model for learning and memory (*37*). We previously showed that the NH125, another small-molecule eEF2K inhibitor that is structurally distinct from AG (*20*), was able to rescue hippocampal LTP deficit induced by exogenous Aβ (*5*). Here, we further demonstrated that application of NH125 (1 μM) on hippocampal slices reversed the LTP impairments displayed in aged APP AD model mice (Fig. 5A-C). LTP performance in hippocampal slices taken from WT mice treated with NH125 was not altered compared to the vehicle-treated group (Fig. 5A-C). These results suggest that pharmacological inhibition of eEF2K via NH125 rescue AD-associated long-term synaptic plasticity deficits.

**Fig. 5.**
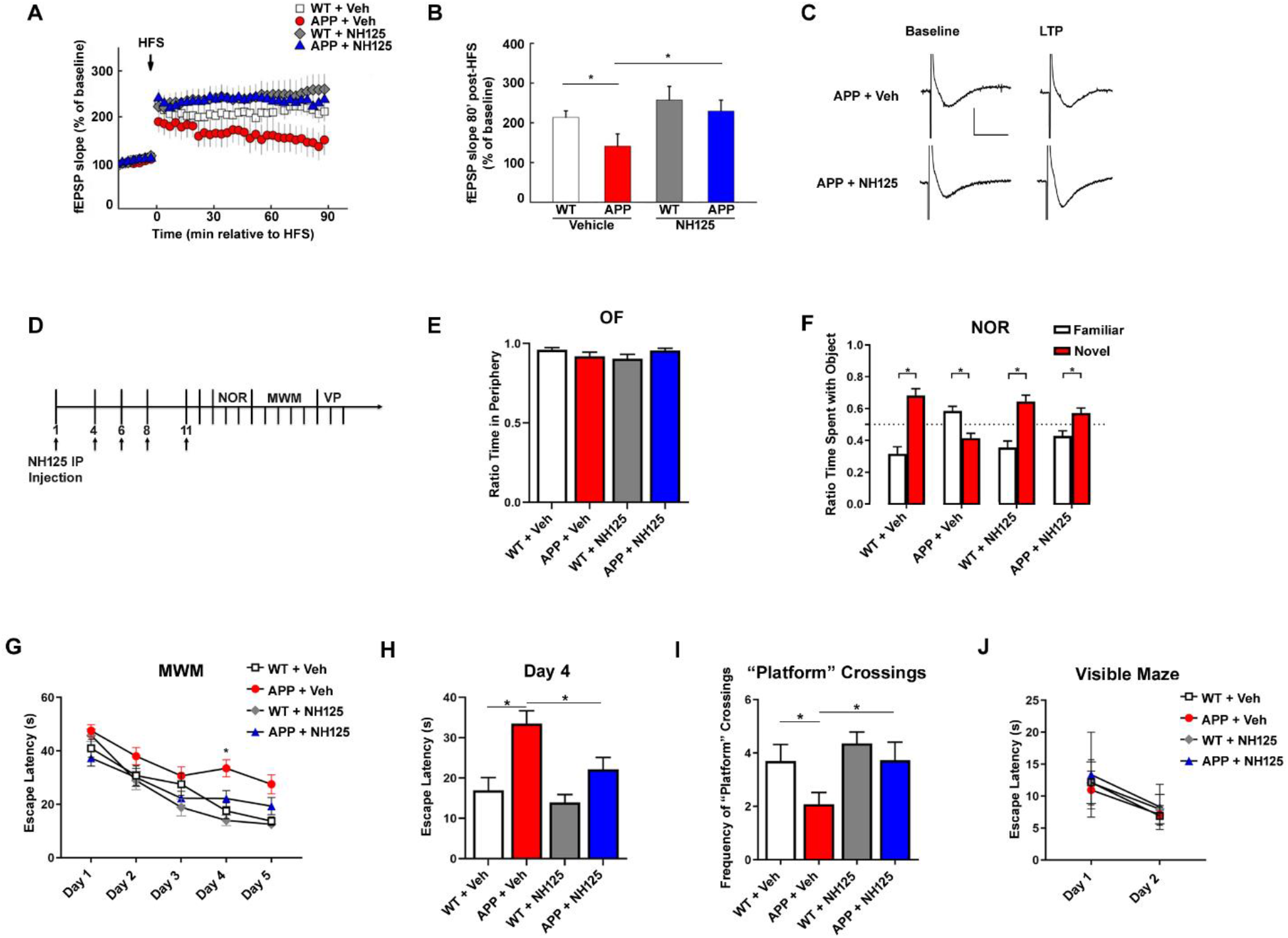
Treatment with eEF2K inhibitor NH125 rescues LTP and cognitive deficits in APP/PS1 AD model mice. **(A)** WT and APP/PS1 slices were treated with vehicle (DMSO; WT, *n* = 8; APP/PS1, *n* = 5) or NH125 (1 μM; WT, *n* = 8; APP/PS1, *n* = 10) and stimulated with HFS to induce LTP. Arrow indicates HFS. APP + Veh slices had significantly impaired LTP compared with WT + Veh, WT + NH125, and APP + NH125 \**p* < 0.05. 1-way ANOVA with Tukey’s post hoc test. **(B)** fEPSP slope 80 minutes after HFS. **(C)** Representative fEPSP traces. **(D)** Treatment schematic showing NH125 injections (1 mg/kg) and subsequent behavioral tasks. **(E)** Ratio of time spent in the periphery for the OF test. (WT + Veh, *n* = 9; APP + Veh, n = 10; WT + NH125, *n* = 8; APP + NH125, *n* = 8). **(F)** Percentage of time spent with familiar (white) and novel (red) objects in the NOR task during the testing phase. Preference of less than 50% indicates cognitive impairment. (WT + Veh, *n* = 9; APP + Veh, *n* = 10; WT + NH125, *n* = 8; APP + NH125, *n* = 8). Statistical preference for novel or familiar object: WT + Veh, \**p* < 0.0001; APP + Veh, *P* = 0.0008; WT + NH125, \**p* = 0.0002; APP + NH125, \**p* = 0.0072, unpaired *t* test. **(G)** Escape latency (s) over 5 days of training in the hidden platform MWM. Four trials/day, 5 days. (WT + Veh, *n* = 13; APP + Veh, *n* = 18; WT + NH125, *n* = 16; APP + NH125, *n* = 16). **(H)** Escape latency (s) on Day 4 of training. WT + Veh versus APP + Veh, \**p* = 0.0062; WT + Veh versus APP + NH125, *p* = 0.5334; APP + Veh versus APP + NH125, \**p* = 0.0002, 1-way ANOVA with Tukey’s post hoc test. *F* = 6.335. **(I)** Frequency of crossing where platform used to be during probe trial phase of MWM task. (WT + Veh, *n* = 13; APP + Veh, *n* = 18; WT + NH125, *n* = 16; APP + NH125, *n* = 16). WT + Veh versus APP + Veh, \**p* = 0.0460; WT + Veh versus APP + NH125, *p* > 0.9999; APP + Veh versus APP + NH125, *\*p* = 0.0309, 1-way ANOVA with Tukey’s post hoc test. *F* = 4.879. **(J)** Escape latency (s) over 2 days in the visible maze task. (WT + Veh, *n* = 13; APP + Veh, *n* = 11; WT + NH125, *n* = 16; APP + NH125, *n* = 12). Error bars represent + SEM.

To further investigate whether pharmacological eEF2K inhibition could alleviate AD-associated cognitive impairments, we administered NH125 (or vehicle control) to aged APP mice followed by a series of behavioral assays to evaluate their cognitive performance. Following previous studies, NH125 (or vehicle) was delivered intermittently through intraperitoneal (IP) injections at a dose of 1 mg/kg (*38*) (Fig. 5D). We observed no significant differences among the four treatment groups in OF performance, including ratio time spent in peripheral/central area of the arena and travel distance and velocity (Fig. 5E, Fig. S4A and B). Next, we performed the NOR task to evaluate long-term recognition memory (*22*). Consistent with the experiments described above (Fig. 1C), the APP mice treated with vehicle spent more time with the familiar object than the novel object, indicating cognitive deficit and behavioral inflexibility (Fig. 5F). Notably, the APP mice treated with NH125 (APP + NH125) spent significantly more time with the novel than with the familiar objects, indicative of normal recognition memory (Fig. 5F).

In the hidden-platform MWM task, WT mice treated with vehicle or NH125 exhibited normal learning and memory as indicated by marked day-to-day decreases in escape latency during the acquisition phase (Fig. 5G and H). APP mice treated with vehicle exhibited longer escape latency times, indicative of a cognitive impairment (Fig. 5G and H). During the probe trial, APP mice crossed the “platform” less frequently compared to WT mice, indicating memory impairments (Fig. 5I). Importantly, impairments of spatial learning and memory associated with APP mice were alleviated by treatment with NH125, as indicated by decreased escape latency and improved “platform” crossings in the probe trial (Fig. 5G-I). Distance traveled and velocity of movement during the probe trial of MWM were not altered across all groups (Fig. S4C and D). Additionally, we examined potential memory-independent effects of NH125 through the visible maze task, and did not find differences in latency to locate the visible platform across all groups (Fig. 5J). Taken together with the findings from the NOR test, these results show that treatment with eEF2K inhibitor NH125 rescues cognitive deficits in APP/PS1 AD model mice.

### Knockout of eEF2K alleviates cognitive deficits in APP/PS1 AD model mice

We further took a genetic approach to completely suppress eEF2K activity in AD by crossing APP/PS1 AD model mice with global homozygous eEF2K knockout mice (eEF2K^−/−^) (see method section for detailed breeding scheme). Four experimental genotype groups were generated including: WT, APP/PS1 (APP), eEF2K^−/−^, and APP/PS1/eEF2K^−/−^ double mutant (APP/eEF2K^−/−^) (Fig. 6A). Both male and female mice aged 12 to 16 months were used for experimentation. Consistent with previous studies (*10, 19*), levels of eEF2 phosphorylation in hippocampal lysates were increased in APP mice compared with WT littermates (Fig. S5A-C). Knockout of eEF2K drastically reduced eEF2 phosphorylation in the WT and APP mice (Fig. S5A-C).

**Fig. 6.**
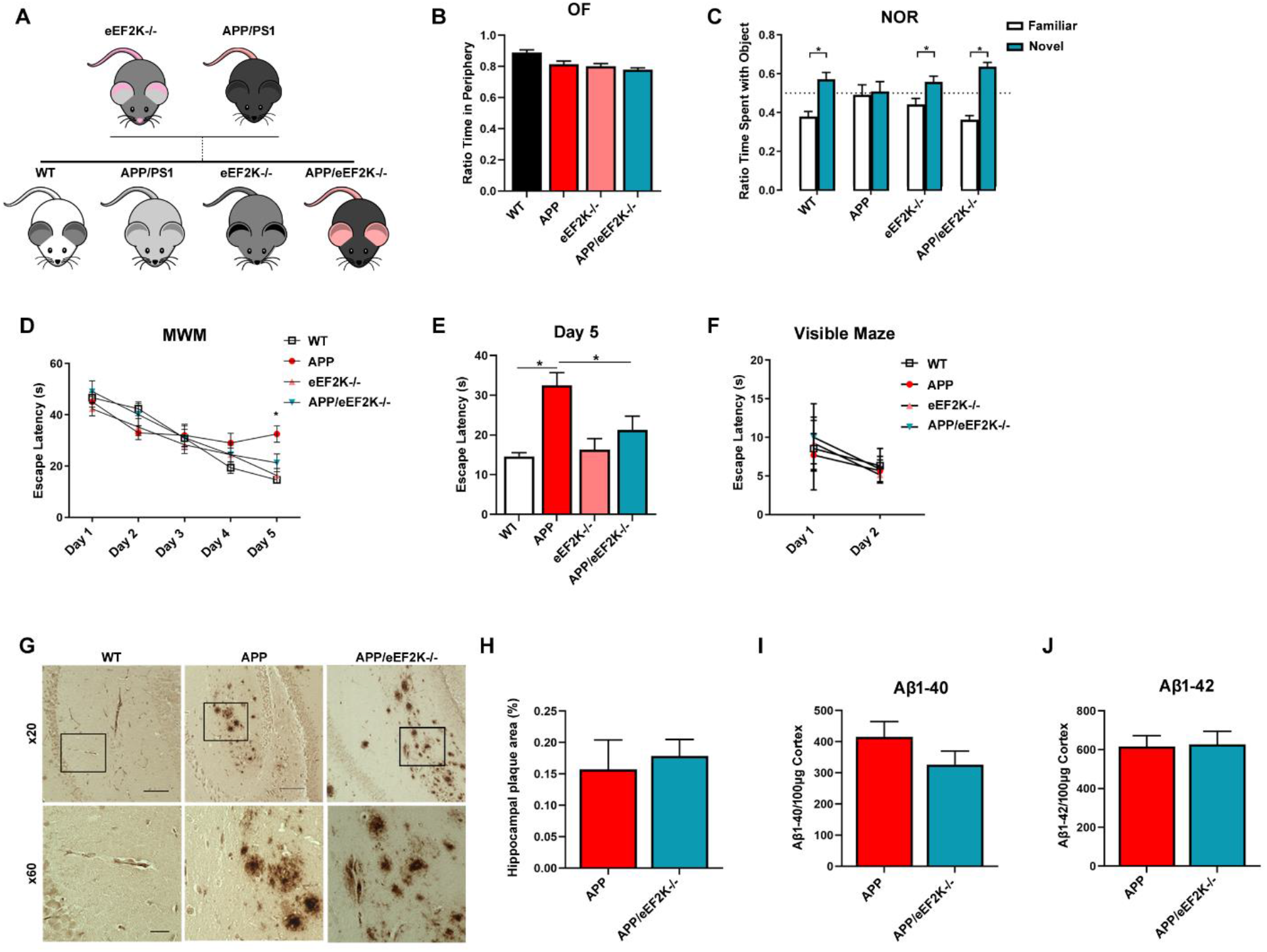
Genetic knockout of eEF2K alleviates AD-associated cognitive deficits. **(A)** Breeding scheme for generating APP/PS1/eEF2K^−/−^ double mutant mice and littermates. **(B)** Ratio of time spent in the periphery during the OF test. (WT, *n* = 17; APP, *n* = 9; eEF2K−/−, *n* = 10; APP/eEF2K−/−, *n* = 8). **(C)** Percentage of time spent with familiar (white) and novel (pink) objects in the NOR task during the testing phase. Preference of less than 50% indicates cognitive impairment. (WT, *n* = 17; APP, *n* = 9; eEF2K−/−, *n* = 10; APP/eEF2K−/−, *n* = 8). Statistical preference for novel or familiar object: WT, \**p* = 0.0002; APP, *p* = 0.822; eEF2K−/−, \**p* = 0.014; APP/eEF2K−/−, \**p* < 0.0001, unpaired *t* tests. **(D)** Escape latency (s) over 5 days of training in the hidden platform MWM. Four trials/day, 5 days. (WT, *n* = 16; APP, *n* = 9; eEF2K−/−, *n* = 16; APP/eEF2K−/−, *n* = 9). WT versus APP, \**p* < 0.0001; APP versus APP/eEF2K Homo, \**p* = 0.04, 1-way ANOVA with Tukey’s *post hoc* test. *F* = 8.888. **(E)** Escape latency (s) on day 5 of MWM training. (WT, *n* = 16; APP, *n* = 9; eEF2K−/−, *n* = 16; APP/eEF2K−/−, *n* = 9). WT versus APP, \**p* < 0.0001; APP versus APP/eEF2K−/−, \**p* = 0.04, 1-way ANOVA with Tukey’s post hoc test. *F* = 8.888. **(F)** Escape latency (s) over 2 days in the visible maze task. (WT, *n* = 16; APP, *n* = 9; eEF2K−/−, *n* = 16; APP/eEF2K−/−, *n* = 9). **(G)** Representative images of hippocampal plaque deposition in WT, APP, and APP/eEF2K−/− mice. Insets are shown at x60 magnification. Scale bars: 100 μm (x20); 20 μm (x60). **(H)** Percentage of hippocampal area covered in amyloid plaques in APP (*n* = 10 sections) and APP/eEF2K−/− (*n* = 8). *p* = 0.70, unpaired *t* test. **(I)** ELISA showed no differences in levels of Aβ 1-40 or Aβ 1-42 **(J)** in APP and APP/eEF2K−/− forebrain tissue. *n* = 6. *p* = 0.22 for Aβ 1-40, *p* = 0.90 for Aβ 1-42, unpaired *t* test. Error bars represent + SEM.

To determine the effects of genetic knockout of eEF2K on AD-associated cognitive impairments, we subjected these mice to the same series of behavioral tasks as described above. In brief, the OF test did not reveal any differences among the 4 genotypes (Fig. 6B, Fig. S5D and E). In the NOR task, recognition memory impairments displayed in APP mice were improved with genetic eEF2K knockout (Fig. 6C). Consistently, deficiency in spatial learning and memory in APP mice, assessed by MWM task, was also alleviated with deletion of the eEF2K gene (Fig. 6D and E). Distance traveled and velocity of movement in the probe trial of MWM were not different across all groups (Fig. S5F and G). Additionally, no difference in latency to locate the visible platform were observed across all groups (Fig. 6F). Taken together, these results show that global eEF2K knockout alleviates AD-associated learning and memory deficits.

We investigated whether genetic knockout of eEF2K would have an effect on AD-associated brain pathology. We did not observe any differences in gross morphology of hippocampus among the 4 genotypes (Fig. S5H). We also assessed Aβ plaque pathology in the hippocampal areas of these mice, and found no significant changes in plaque deposition in APP/eEF2K^−/−^ mice, compared to APP group (Fig. 6G and H). We then assessed brain levels of Aβ 1-42 and Aβ 1-40 using ELISA, and found no significant differences between APP and APP/eEF2K^−/−^ mice (Fig. 6I and J). Further, we assessed levels of phosphorylated tau using a biochemical approach, and did not find significant differences in the levels of tau phosphorylation across all 4 genotypes (Fig. S5I-K).

## Discussion

It is urgent to identify alternative therapeutic strategies for AD based on thorough mechanistic studies. In the current study, we have shown that suppression of eEF2K activity with two structurally distinct small-molecule inhibitors rescued aging-related cognitive deficits in two different strains of AD model mice. Our studies demonstrated that systemic application of small-molecule inhibitors of eEF2K rescues multiple pathologies characteristic of AD. Together with our studies using genetic approaches, these findings support the concept that eEF2K suppression and perhaps restoration of overall translational elongation capacity could be a feasible therapeutic strategy for AD and related dementia syndromes.

Mounting evidence indicates a role of eEF2K in human diseases including solid cancer, cardiovascular diseases, and neuronal disorders (*39*). Interestingly, both NH125 and A-484954 were originally developed for cancer treatment (*20, 39, 40*). From a perspective of translational medicine, several attractive features of eEF2K could make it an ideal target of intervention for AD treatment. First, mounting evidence demonstrates that eEF2K activity is not required for development or cell survival under physiological conditions. Mice with global knockout of the eEF2K gene appear normal during different stages of development as demonstrated in multiple studies (*19, 41, 42*). We also did not observe any adverse effects in WT mice treated with the eEF2K inhibitor AG, including weight changes and gross morphology of hippocampus (Fig. S1C and H-I). Thus, it is promising for future studies in a clinical setting that eEF2K inhibitors can be given safely without causing serious side effects. This (the safety issue) is critically important for AD patients who usually need to keep taking medicine over a long period of time. Second, eEF2K belongs to a small group of kinases termed “alpha-kinases”, whose catalytic domains are distinct from those in the conventional protein kinases (e.g. serine/threonine kinases and tyrosine kinases), which consists of the vast majority of the eukaryotic protein kinase superfamily (*43*). Therefore, compounds targeting eEF2K are unlikely to affect activities of other conventional kinases that play important roles for a broad spectrum of biological processes. Third, eEF2K is the only known kinase for eEF2 and eEF2 is the only known substrate for eEF2K (phosphorylation at the Thr56 site) (*16, 17*). This one-to-one relationship can further strengthen the specificity and selectivity of the eEF2K antagonists if they are properly designed for therapeutic use.

What are the potential molecular mechanisms underlying the rescue effects of the eEF2K inhibitors on AD-related cognitive impairments? Repression of eEF2K leads to de-phosphorylation of eEF2 and consequently increases overall protein synthesis (*12*). Impaired protein synthesis has been demonstrated in AD brains (*16, 44*). Thus, the parsimonious explanation would be improvement of protein synthesis, which raises another significant question: are the rescued phenotypes by the eEF2K inhibitors associated with alterations or restoration of any specific proteins controlled by eEF2K/eEF2 signaling? As a starting point to address this question, we used an unbiased proteomics approach and identified multiple proteins whose levels are dysregulated (increased or decreased) in the brain of AD mice and restored with treatment of the eEF2K inhibitor AG (Fig. 3E-G). Interestingly, for most (8 out of 11) of the identified proteins based on the aforementioned criteria, the protein expression levels are increased in the brain of AD mice and reduced with eEF2K inhibition (Fig. 3E-G). Such findings seem counterintuitive given that overall protein synthesis capacity is decreased in AD mice and that inhibition of eEF2K improves general protein synthesis (Fig. 3A-D). Future in-depth studies including those at the functional levels shall help determine the association of the dysregulation of these proteins and AD pathogenesis, and their relationship (e.g. direct or indirect substrates) with eEF2K/eEF2 signaling. Moreover, decades of research has not yet reached a consensus with regards to the identities of “memory proteins” or “plasticity-related proteins (PRPs)” (*45*). It is reasonable to speculate that, instead of regulation on specific proteins (e.g. PRPs), an improvement of “general” protein synthesis through a boost of translational capacity (e.g. eEF2K inhibition) can account for alleviation of AD-associated cognitive deficits and synaptic failure.

It needs to be cautioned that previous studies conducted in non-neuronal cell lines argued the specificity and inhibition potency of NH125 on eEF2K, as well as the actual effects on eEF2 phosphorylation (*20, 46*). Elucidation of potential eEF2K-independent mechanisms underlying the rescuing effects of NH125 on AD-associated synaptic failure and cognitive impairments warrants further in-depth studies. There are other caveats to be considered before translation of this strategy into human patients. We studied the effects of the eEF2K inhibitors in two mouse models of AD, both are models for familial Alzheimer’s disease (FAD), which accounts for <5% of all AD cases. We anticipate to validate our findings with eEF2K inhibition in sporadic AD (SAD) models once they are established. We observed that the treatment with eEF2K inhibitors does not alter brain Aβ pathology, which is consistent with our previous studies using a genetic approach to suppress eEF2K in AD models (*19*). Meanwhile, pharmacological inhibition of eEF2K in both models significantly improved cognitive impairments, synapse loss, and protein synthesis defects, which are exemplified in both FAD and SAD. Moreover, we are glad to see, from a therapeutic point of view, that application of the eEF2K inhibitors is effective at the late, symptomatic stage of the AD model mice. It is also worth mentioning that activation of eEF2K and increased eEF2 phosphorylation may function as a beneficial cellular stress response in situations such as nutrient starvation or hypoxia (*12*). Thus, it is possible that hyper-phosphorylation of eEF2 and protein synthesis inhibition are actually neuroprotective at the very early stage of AD. It would be informative in the future to characterize the effects of eEF2K intervention at different stages of AD pathogenesis to provide insights for more accurate design of clinical studies.

## Materials and Methods

### Study Design

We utilized a power analysis to calculate the sample size necessary to achieve a reliable measurement of the effect, and we were able to utilize the appropriate number of animals according to the preliminary power analysis. We used a Grubbs test in order to identify outliers in all data sets, and if outliers were identified, those data points were excluded. The objectives of this study were to test the potential for eEF2K inhibitors as a treatment for AD. Our hypothesis was that administration of eEF2K inhibitors NH125 or AG would alleviate cognitive deficits in AD model mice, and that protein synthesis capacity would be restored. Once that hypothesis was proven, we then hypothesized that AD model mice treated with AG would demonstrate rescued spine morphology and density, post-synaptic density formation, and polyribosome formation, and that these improvements could occur independently of amyloid beta and tau pathology. The research subjects were animals, specifically APP/PS1 and Tg19959 AD model mice. This study was a controlled laboratory experiment, and mice were randomly assigned to their treatment group. The study was blinded, and the experimenters did not know the allocated condition of the animals when performing behavioral experiments or analyzing tissue.

### Mice

All mice were housed at the Wake Forest School of Medicine barrier facility under the supervision of the Animal Research Program. Mice adhered to a 12-hour light/12-hour dark cycle, with regular feeding, cage cleaning, and 24-hour food and water access. Both male and female mice were used for experimentation. APP/PS1 mice were purchased from the Jackson Laboratory and expressed human transgenes for APP (KM670/671NL) and presenilin-1 (PSEN1 L166P) (*47*). APP/PS1 mice were crossbred with eEF2K^−/−^ mice to first generate APP/PS1/eEF2K^+/−^ double-mutant mice, which were then crossed with eEF2K^+/−^ mice to generate the APP/PS1/eEF2K^−/−^ double-mutant mice (APP/eEF2K^−/−^) and other experimental groups including WT, APP/PS1, and eEF2K^−/−^. All genotyping was done by polymerase chain reaction (PCR). Briefly, Tg19959 mice overexpress mutant human amyloid precursor proteins (APP KM670/671NL, V717F) (*23*). Breeders of Tg19959 AD model mice were a gift from Dr. George Carlson of McLaughlin Research Institute (Great Falls, Montana, USA). Tg19959 cohorts underwent all experimental protocols at 6 to 9 months of age, while APP/PS1 cohorts were evaluated at 12 to 16 months of age (*23, 47*).

### A-484954 treatment

A-484954 (Millipore, catalog 324516-10MG) was sent to Innovative Research of America (Sarasota, Florida), where pellets were manufactured. Pellets were stored at room temperature. Mice were anesthetized using the “SomnoSuite” anesthesia system (Kent Scientific), and sedation of mice was measured with a paw pinch. Once mice were adequately sedated, a pellet containing either A-484954 or vehicle was placed into a 10 gauge trochar. Pellets contained 2.625 mg, a dose previously established to induce effects on eEF2K in mice (*21*), of either A-484954 or vehicle. The pellet dissolved over 30 days. The skin was pierced with the trochar and the pellet was placed subcutaneously. Following placement, mice were monitored for negative side-effects of the drug and to ensure lack of injury from surgery. Two weeks after placement, behavior tasks began.

### Mouse behavioral assays

#### Open field (OF)

Mice were handled for at least 5 days prior to behavioral testing and habituated to the testing facility for an hour prior to experimentation. Animals were placed in an opaque plastic OF chamber (40 cm x 40 cm x 40 cm) and allowed to explore for 15 minutes. Time spent in the center and periphery of the chamber was measured and calculated as a percentage of total time. Distance moved and velocity were measured using EthoVision XT Tracking Software (Noldus Information Technology). Data collection and analysis were performed blinded.

#### Novel object recognition (NOR)

Mice underwent a 2-day familiarization protocol in which they were placed in an opaque, plastic arena (40 cm x 40 cm x 40 cm) with 2 identical objects and allowed to explore for 5 minutes. Twenty-four hours after familiarization, animals were tested in the arena for 5 minutes with one object replaced with a novel object. All objects were randomly assigned to mice, and placement of novel objects was counterbalanced. Time spent with each object was measured and calculated as a percentage of the total interaction time. Novel object preference of less than 50% indicates memory impairment. Time with objects was measured both manually and using EthoVision XT Tracking Software. Mice with a total interaction time of less than 10 seconds were excluded from analysis. Data collection and analysis were performed blinded.

#### Morris water maze (MWM)

MWM was performed as previously described (*8, 19*). The paradigm consisted of 4 trials (60-second maximum, 15-minute interval) per day for 5 consecutive days. Escape latency was measured each training day. A probe trial was performed 2 hours following training on the fifth day. The visible maze task consisted of 4 trials per day for 2 consecutive days, with the escape platform marked by a visible cue and moved randomly among 4 locations. Trajectories, time spent in maze quadrant, velocity, and distance moved were recorded using EthoVision XT Tracking Software. Data collection and analysis were performed blinded.

#### Western blot assay

Mouse hippocampal tissue was flash-frozen on dry ice and sonicated as previously described in lysis buffer with protease and phosphatase inhibitors (*48*). Samples containing equal amounts of protein lysate were loaded on 4%-12% Tris-glycine SDS-PAGE (Bio-Rad) gels for standard gel electrophoresis. Following transfer, nitrocellulose membranes were blocked for 10 minutes in SuperBlock TBS Blocking Buffer (Thermo Fischer Scientific). All primary and secondary antibodies were diluted in 5% milk/TBST or 5% BSA/TBST. Blots were probed with primary antibodies for phospho-eEF2 (Thr56) (1:1000, Cell Signaling Technology, catalog 2331), eEF2 (1:1000, Cell Signaling Technology, catalog 2332), phospho-tau (Ser396) (1:1000, Thermo Fischer, catalog 44-752G), tau (1:1000, Sigma-Aldrich, catalog T5530), phospho-p70 S6 Kinase (1:1000, Cell Signaling Technology, catalog CS9204), p70 (1:1000, Cell Signal, catalog 2708), phospho-eIF2a (Ser51) (1:1000, Cell Signaling Technology, catalog CS3398), eIF2a (1:1000, Cell Signaling Technology, catalog CS9722), phospho-AMPKa (Thr172) (1:1000, Cell Signaling Technology, catalog CS2535), AMPKa (1:1000, Cell Signaling Technology, catalog CS5832), phospho-mTOR (Ser2448) (1:1000, Cell Signaling Technology, catalog CS5536), phospho-mTOR 2481 (1:1000, Cell Signaling Technology, catalog CS2974), mTOR (1:1000, Cell Signaling Technology, catalog CS2983), β-actin (1:10,000, MilliporeSigma, catalog A2228), GAPDH (1:10,000, Cell Signal, catalog 2118). Proteins were visualized using the ChemiDoc Imaging System (Bio-Rad). Densitometric analysis was performed using ImageJ software (NIH).

#### High Performance Liquid Chromatography (HPLC)

Tissue samples were weighed into homogenization tubes (Thermo Fisher Scientific, Waltham, MA, USA) containing 1.4 mm ceramic beads and homogenized after the addition of 500 μL of methanol (Optima, Thermo Fisher Scientific, Waltham, MA, USA) using a Bead Ruptor 24 (OMNI International, Kennesaw, Georgia). The homogenates were then centrifuged at 16,000 x g for 15 minutes. The resulting supernatant was removed and dried under vacuum before being reconstituted in 5% acetonitrile in water (Optima, Thermo Fisher Scientific, Waltham, MA, USA) for LC-MS/MS analysis.

Samples were analyzed using a Shimadzu Nexera UHPLC coupled with a Shimadzu LCMS 8050 triple quadrupole mass spectrometer (Shimadzu, Kyoto, Japan). Ionization in the DUIS source used the following parameters: nebulizing gas flow of 2 L/min, heating gas flow of 10 L/min, interface temperature of 350°C, DL temperature of 200°C, heat block temperature of 400°C, and a drying gas flow of 10 L/min. A-484954 was monitored using the following MRM transitions: 290.10 > 273.10, 202.10, 219.10. The 290.10 > 271.10 transition was used for analyte quantification and the other transitions were monitored as references. Samples were quantified using a standard curve created using a genuine standard. LabSolutions data review software (Version 5.72; Shimadzu, Kyoto, Japan) was used for processing data.

A-484954 was retained on a Zorbax C18 (100 x 2.1 mm, 1.8 μm; Agilent, Santa Clara, CA USA) and eluted with a flow rate of 0.4 ml/minute and mobile phases consisting of 0.1% formic acid for mobile phase A and acetonitrile for mobile phase B. The mobile phase gradient began at 5% B and held there for 0.75 minutes. The gradient was then increased to 95% B at 6.0 minutes. This percentage was held for 1.5 minutes before being decreased to 5% B at 8.0 minutes and then held at that final percentage until 10.0 minutes.

#### Mouse tissue immunohistochemistry

Following euthanasia, mouse brains were hemisected and fixed overnight in ice-cold PFA and transferred to 70% EtOH. Paraffin embedding was performed by Wake Forest Pathology core service. Paraffin-embedded sections (5 μm) mounted on charged slides were cleared in xylene and rehydrated in a graded ethanol series. Antigen retrieval used citrate buffer (pH 6.0) in a standard 10-minute microwave procedure. Blocking was done for 2 hours with 10% NGS in 1% BSA/TBS. Slides were incubated in a humidified chamber in primary antibody for amyloid-β (6E10) (1:200) overnight at 4°C. Following a 15-minute blocking in 3% hydrogen peroxide, sections were incubated in biotinylated anti-mouse secondary antibody (1:200, Vector Labs) for 1 hour at room temperature, followed by Vectastain Elite ABC Reagent (Vector Labs) for another 30 minutes. Primary and secondary antibodies as well as ABC reagent were diluted in 1% BSA/TBS. Sections were developed in DAB (Vector Labs) for 30 seconds to 3 minutes with monitoring. Slides were counterstained using Mayer’s hematoxylin for 60 seconds and stained blue with 0.2% lithium carbonate for 20 seconds. In between each step of immunohistochemistry, sections were rinsed using distilled water or TBSTX (pH 7.4). Negative controls were incubated in 1% BSA with no primary antibody. Sections were dehydrated in an alcohol series and cleared with xylene, coverslipped, and dried overnight. Imaging was performed using BZ-X710 All-in-One Fluorescent Microscope (Keyence).

#### Aβ ELISA

Frozen mouse forebrain samples were sonicated as previously described (*10*). Samples were centrifuged at 16,000 *g* for 20 minutes at 4°C. The supernatant was collected for ELISA. Aβ 1-42 (Thermo Fischer Scientific, catalog KMB3441) and Aβ 1-40 (Thermo Fischer Scientific, catalog KMB3481). ELISAs were performed according to the manufacturer’s instructions. Ninety-six-well plates were read at 450 nm using an iMark microplate reader (Bio-Rad).

#### SUnSET assay

Acute 400 μm transverse hippocampal slices were prepared using a Leica VT1200S vibratome as described previously (*10*). Slices were maintained before experimentation at room temperature for at least 2 hours in ACSF containing the following (in mM): 118 NaCl, 3.5 KCl, 2.5 CaCl_2_, 1.3 MgSO_4_, 1.25 NaH_2_PO_4_, and 15 glucose, bubbled with 95% O_2_ / 5% CO_2_. Slices were incubated in puromycin (1 μg/ml) for 1 hour at 32°C in bubbling ACSF. Slices were then flash-frozen on dry ice, and area CA1 was microdissected for Western blot analysis. Puromycin-labeled proteins were identified using the mouse monoclonal antibody (1:10,000, Millipore, catalog MABE343). Protein synthesis levels were determined by analyzing total lane density from 10 kDa to 250 kDa. Densitometric analysis was performed using ImageJ software (NIH).

#### Mass spectrometry-based proteomic analysis

Whole prefrontal cortices were flash frozen on dry ice, dissected in ice-cold PBS, and lysed in 500 μL of PBS with protease/phosphatase inhibitor using a Bead Mill Homogenizer (Bead Ruptor, Omni International); 500 μL of x2 RIPA buffer was added, and the mixture was incubated on ice for 30 minutes before centrifugation at 18,000 *g* for 10 minutes. Protein amount was measured in supernatant, and 50 μg of protein was subjected to tryptic digestion.

Reducing alkylation was performed in the presence of 10 mM dithiothreitol and 30 mM iodoacetamide. Four times the sample volume of cold acetone was added to the tube, which was incubated at −20°C overnight. Tubes were centrifuged at 14,000 *g* for 10 minutes to obtain pellet, which was resuspended in 50 mM ammonium bicarbonate; 1 μg of sequencing-grade modified trypsin was added (1:50 enzyme to substrate) and incubated at 37°C overnight. Tryptic digest was purified using a C18 desalting spin column and then prepared in 5% (v/v) CAN containing 1% (v/v) formic acid for liquid chromatography-tandem MS (LC-MS/MS) analysis.

The LC-MS/MS system consisted of Q Exactive HF Hybrid Quadruple-Orbitrap Mass Spectrometer (Thermo Fischer Scientific) and a Dionex Ultimate-3000 Nano-UPLC System (Thermo Fischer Scientific) employing a Nanospray Flex Ion Source (Thermo Fischer Scientific). An Acclaim PepMap 100 (C18, 5 μm, 100 Å, 100 μm x 2 cm) trap column and an Acclaim PepMap RSLC (C18, 2 μm, 100 Å, 75 μm x 50 cm) analytical column were used for the stationary phase. Peptides were separated employing a linear gradient consisting of mobile phases A (water with 0.1% formic acid) and B (acetonitrile with 0.1% formic acid), where the gradient was from 5% B at 0 minutes to 40% B at 170 minutes. MS spectra were acquired by data-dependent scans consisting of MS/MS scans of the 20 most intense ions from the full MS scan with a dynamic exclusion option, which was 10 seconds.

Spectra were searched using the Sequest HT algorithm within the Proteome Discoverer, version 2.1 (Thermo Fischer Scientific), in combination with the mouse UniProt protein FASTA database (annotated 16,747 entries, December 2015). Search parameters were as follows; FT-trap instrument, parent mass error tolerance of 10 ppm, fragment mass error tolerance of 0.02 Da (monoisotopic), variable modifications of 16 Da (oxidation) on methionine, and fixed modification of 57 Da (carbamidomethylation) on cysteine. Normalized peak intensity data were analyzed for all samples. Using R coding software, 1-way ANOVA was performed for each protein reported, and those with general significance were further investigated. Tukey’s *post hoc* tests were performed, and the final proteins chosen had to meet two criteria: (1) significant difference between WT + Veh and Tg + Veh and (2) significant difference between Tg + Veh and Tg + AG. The proteins that matched these criteria were then entered into the UniProt online database (https://www.uniprot.org/) to determine functional category of each protein. Pie charts were created using RapidTables online software (https://www.rapidtables.com/tools/pie-chart.html).

#### Transmission electron microscopy (TEM)

Brains were removed, and 1 mm thick transverse slices were cut using a Leica VT1200S vibratome. The CA1 was dissected and immediately fixed in 2.5% glutaraldehyde/1% paraformaldehyde in 0.1M Millonig’s phosphate buffer (pH 7.3) overnight. The samples were washed in buffer and post-fixed with 1% osmium tetroxide in phosphate buffer for 1 hour. After washing, samples were dehydrated through a graded series of ethanol solutions. For preparation of resin infiltration, the samples were incubated in propylene oxide for two 15-minute changes. Finally, the samples were gradually infiltrated with 1:1, 1:2, and pure solutions of Spurr’s resin and cured in a 70°C oven overnight. Sections of 90 nm were obtained with a Reichert-Jung Ultra-cut E ultramicrotome, stained with lead citrate and uranyl acetate, and viewed with a Tecnai Spirit transmission electron microscope operating at 80 kV (FEI Co.). Images were obtained with a 2Vu CCD camera (Advanced Microscopy Techniques) at 11,000. Analysis for PSDs and polyribosomes was performed as previously described (*19*). Imaging and analysis were done blinded to animal groups.

#### Golgi-Cox stain and spine morphology analysis

Brains were processed using the FD Rapid GolgiStain Kit in accordance with the manufacturer’s instructions (FD Neurotechnologies, catalog PK401). Transverse sections (100 μm) were made using a Leica VT1200S vibratome and mounted onto gelatin-coated slides. Development was performed according to kit instructions. Sections were dehydrated through a graded ethanol series and cleared in xylene. Slides were coverslipped with Vecta-Mount Permanent Mounting Medium (Vector Labs, catalog H-5000) and imaged at x100 on a Keyence BZ-X710 microscope. Area CA1 stratum radiatum apical dendrites were quantified. For spine analysis, images were blinded, and spines were manually counted and sorted as previously described (*34*).

#### Hippocampal slice preparation and electrophysiology

Acute 400 μm transverse hippocampal slices were prepared using a Leica VT1200S vibratome, as described previously (*48*). Slices were maintained before experimentation at room temperature for at least 2 hours in artificial cerebrospinal fluid (ACSF) containing the following (in mM): 118 NaCl, 3.5 KCl, 2.5 CaCl_2_, 1.3 MgSO_4_, 1.25 NaH_2_PO_4_, and 15 glucose, bubbled with 95% O_2_ / 5% CO_2_. For electrophysiology, monophasic, constant-current stimuli (100 μs) was delivered with a bipolar silver electrode placed in the stratum radiatum of area CA3. Field excitatory post-synaptic potentials (fEPSPs) was recorded using a glass microelectrode from the stratum radiatum of area CA1. Long-term potentiation (LTP) was induced using high-frequency stimulation consisting of two 1-sec 100 Hz trains separated by 60 sec, each delivered at 70-80% of the intensity that evoked spiked fEPSPs. Data collection and analysis were not performed blinded.

#### NH125 treatment

NH125 (Millipore, catalog 324515-5MG) stock solution was prepared in DMSO to its final concentration of 1 M before use. NH125 was stored at −20 °C. For injections, stock NH125 was diluted in PBS to a final concentration of 1 mM. Mice were weighed before injection to ensure a dosage of 1 mg/kg. Mice were injected intraperitoneally with either NH125 or PBS with an equivalent amount of DMSO on an established dosing schedule (*38*) over 11 days as follows: Day 1, Day 4, Day 6, Day 8, and Day 11. Mice were monitored for negative side-effects of the drug and to ensure lack of injury from injections.

#### Statistical analysis

Data are presented as a mean + SEM. Summary data are presented as group means with SE bars. For comparisons between 2 groups, a 2-tailed independent Student’s *t* test was performed using Prism 9 software (GraphPad Software). Two-tailed paired *t* tests were performed for within-group analyses. For comparisons among more than 2 groups, 1-way ANOVA was used with Tukey’s post hoc tests for multiple comparisons. Error probabilities of *p* < 0.05 were considered statistically significant unless otherwise noted.

#### Study approval

All protocols involving animals were approved by the Institutional Animal Care and Use Committee of Wake Forest University School of Medicine. Mice were kept in compliance with the NIH *Guide for the Care and Use of Laboratory Animals* (National Academies Press, 2011).

## Supplementary Materials

**Fig. S1.**
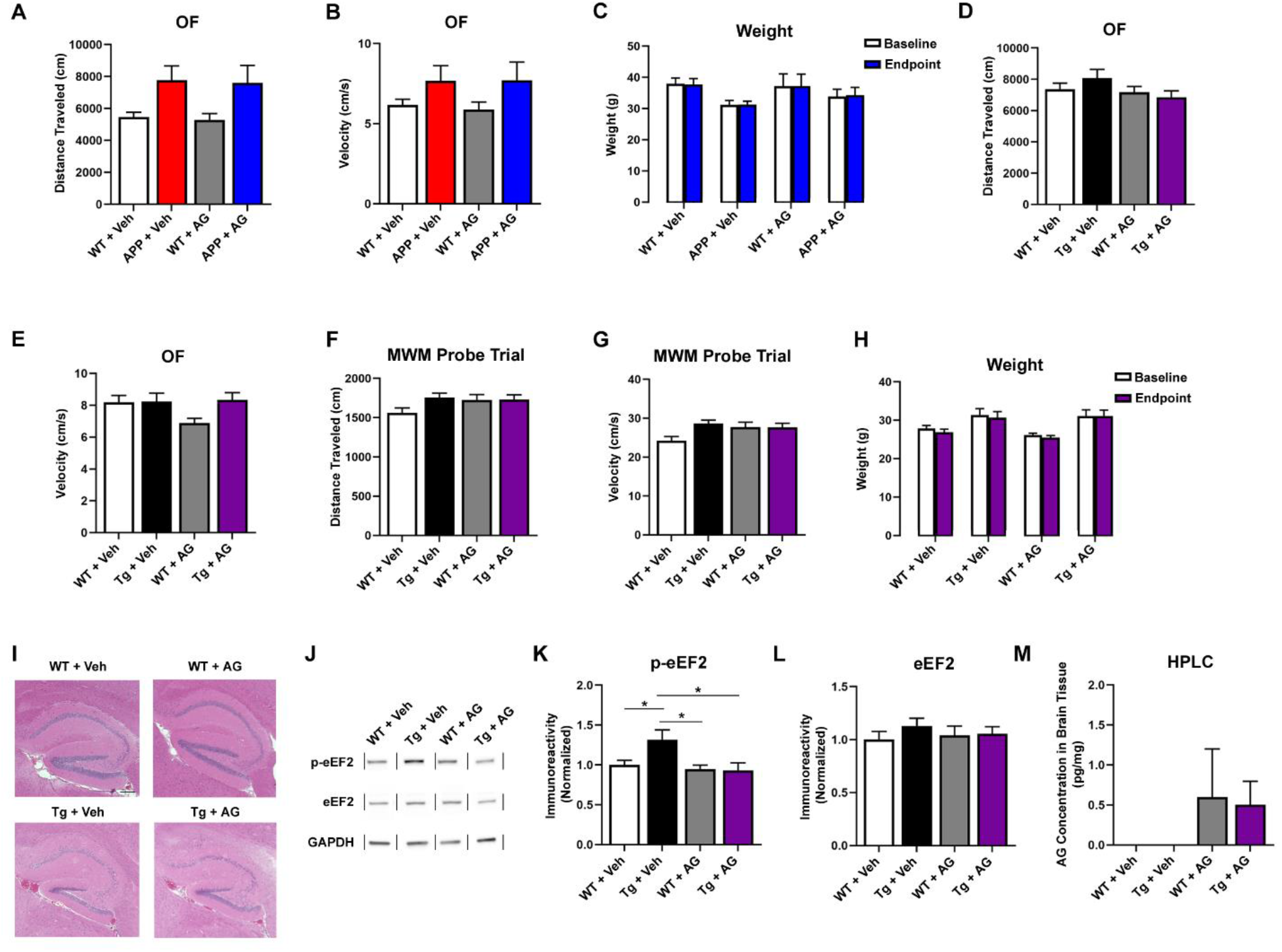
Characterization of phenotypes in AD model mice treated with AG. **(A)** Distance traveled in cm of mice in open field assay (WT + Veh, *n* = 12; APP + Veh, *n* = 6; WT + AG, *n* = 8; APP + AG, *n* = 7). **(B)** Velocity of movement in cm/s for open field assay. **(C)** Baseline and endpoint weight in grams for WT + Veh (*n* = 12), APP + Veh (*n* = 5), WT + AG (*n* = 7), and APP + AG (*n* = 7) experimental mice. **(D)** Distance traveled in cm of mice in open field assay (WT + Veh, *n* = 11; Tg + Veh, *n* = 10; WT + AG, *n* = 13; Tg + AG, *n* = 10). **(E)** Velocity of movement in cm/s for open field assay. **(F)** Distance traveled in cm for MWM probe trial assay (WT + Veh, *n* = 13; Tg + Veh, *n* = 11; WT + AG, *n* = 16; Tg + AG, *n* = 12). **(G)** Velocity of movement in cm/s for MWM probe trial. **(H)** Baseline and endpoint weight in grams for WT + Veh (*n* = 17), Tg + Veh (*n* = 14), WT + AG (*n* = 20), and Tg + AG (*n* = 15) experimental mice. **(I)** Representative images showing hematoxylin & eosin staining in hippocampal sections (20X, scale bar 200 μm). **(J)** Representative Western blot showing p-eEF2 and eEF2 levels in hippocampal lysates. **(K)** Levels of phosphorylated eEF2 were significantly higher in Tg + Veh than WT + Veh lysates, but were significantly lower in Tg + AG than Tg + Veh lysates. (WT + Veh, *n* = 6; Tg + Veh, *n* = 5; WT + AG, *n* = 4; Tg + AG, *n* = 4). WT + Veh versus Tg + Veh, \**p* = 0.03; Tg + Veh versus WT + AG, \**p* = 0.03; Tg + Veh versus Tg + AG, \**p* = 0.03, 1-way ANOVA with Tukey’s post hoc test. *F* = 4.257. **(L)** No difference in the levels of total eEF2 were detected in all groups. **(M)** AG concentration in the brain (pg AG/mg tissue) measured by HPLC (WT + Veh, n = 4; Tg + Veh, n = 4; WT + AG, n = 3; Tg + AG, n = 4).

**Fig. S2.**
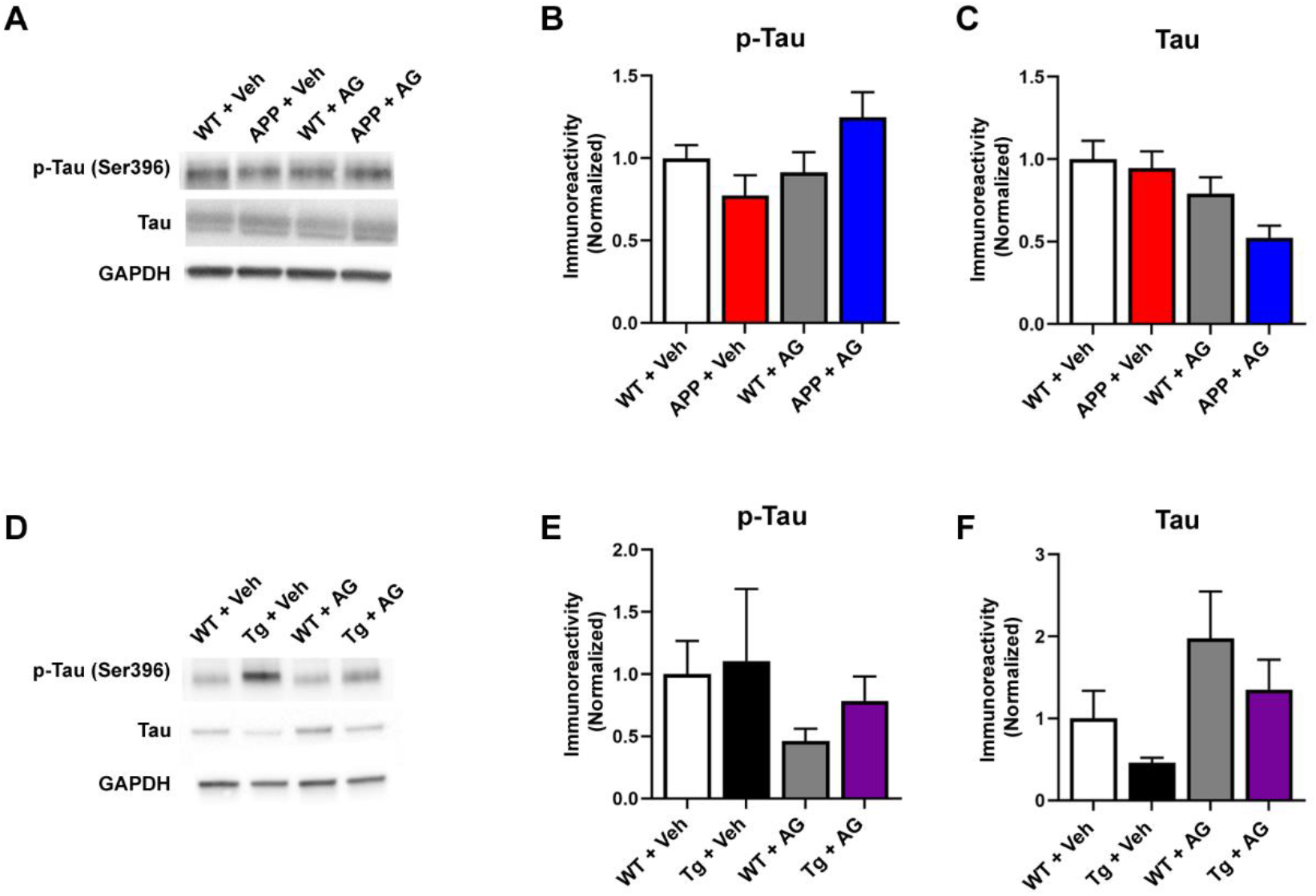
Levels of tau phosphorylation are not affected by treatment of eEF2K inhibitor AG. **(A)** Representative Western blots showing levels of phosphorylated tau and tau in hippocampal lysates. **(B)** No significant differences in levels of phosphorylated tau were detected in WT + Veh (*n* = 5), Tg + Veh (*n* = 5), WT + AG (*n* = 6), and Tg + AG (*n* = 8) experimental mice. **(C)** No significant differences in levels of total tau were detected. **(D)** Representative Western blots showing levels of phosphorylated tau and total tau in hippocampal lysates. **(E)** No significant differences in levels of phosphorylated tau were detected in WT + Veh (*n* = 8), APP + Veh (*n* = 6), WT + AG (*n* = 5), and APP + AG (*n* = 7) experimental mice. **(F)** No significant differences in levels of total tau were detected.

**Fig. S3.**
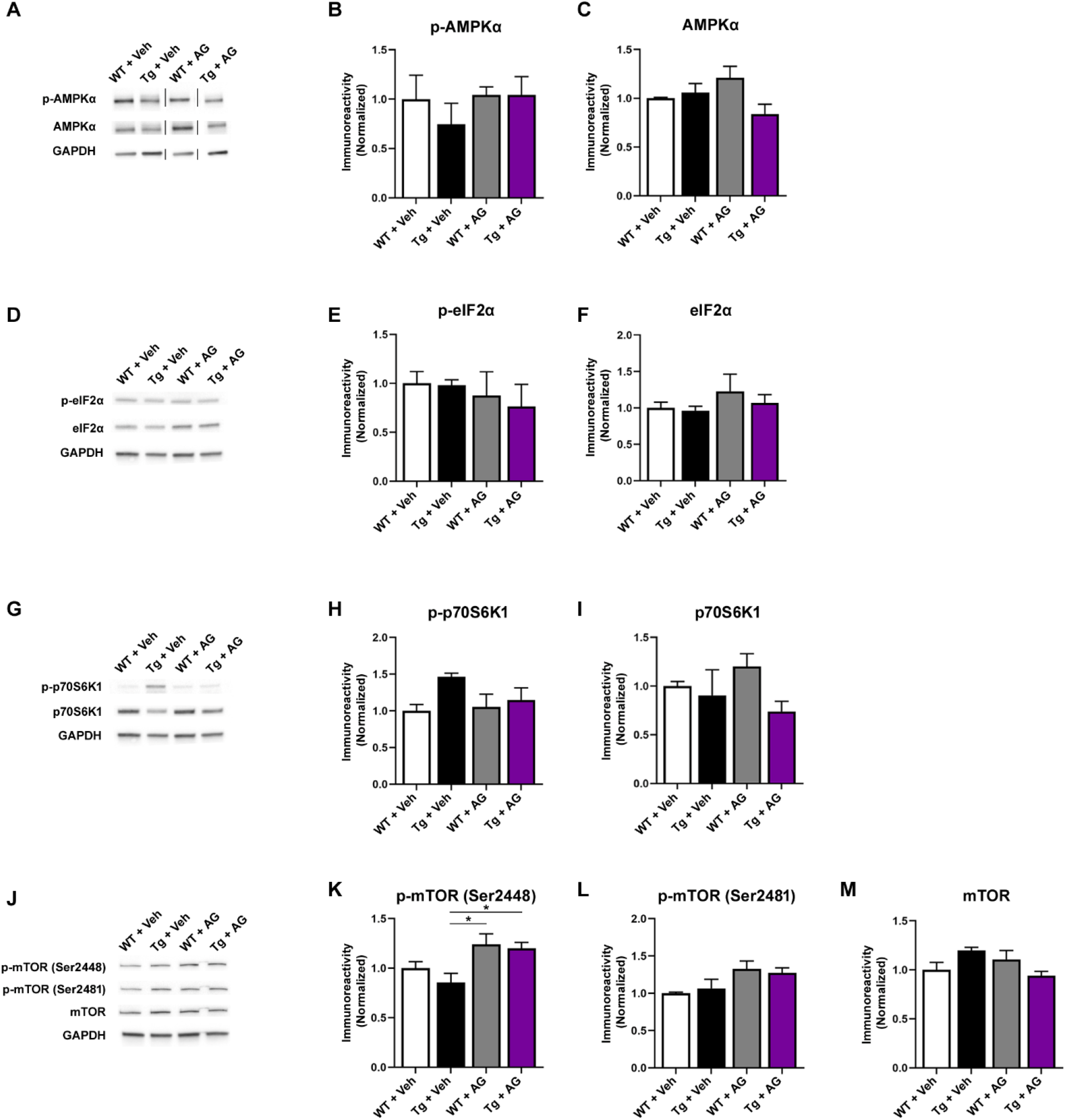
Characterization of signaling pathways related to eEF2K in mice treated with the eEF2K inhibitor AG. **(A)** Representative Western blots showing levels of p-AMPKα, AMPKα, and GAPDH in WT + Veh (*n* = 4), Tg + Veh (*n* = 4), WT + AG (*n* = 6), and Tg + AG (*n* = 5) mice. **(B)** No significant differences in levels of p-AMPKα were detected. **(C)** No significant differences in levels of AMPKα were detected. **(D)** Representative Western blots showing levels of p-eIF2α, eIF2α, and GAPDH in WT + Veh (*n* = 4), Tg + Veh (*n* = 4), WT + AG (*n* = 6), and Tg + AG (*n* = 5) mice. **(E)** No significant differences in levels of p-eIF2α were detected. **(F)** No significant differences in levels of eIF2a were detected. **(G)** Representative Western blots showing levels of p-p70S6K, p70S6K, and GAPDH in WT + Veh (*n* = 4), Tg + Veh (*n* = 4), WT + AG (*n* = 6), and Tg + AG (*n* = 5) mice. **(H)** No significant differences in levels of p-p70S6K were detected. **(I)** No significant differences in levels of p70S6K were detected. **(J)** Representative Western blots showing levels of p-mTOR (Ser2448), p-mTOR (Ser2481), mTOR, and GAPDH in WT + Veh (*n* = 4), Tg + Veh (*n* = 4), WT + AG (*n* = 6), and Tg + AG (*n* = 5) mice. **(K)** Levels of p-mTOR (Ser2448) were significantly lower in Tg + Veh mice compared to WT + AG and Tg + AG mice (Tg + Veh versus WT + AG, \**p* = 0.03; Tg + Veh versus Tg + AG, \**p* = 0.04, 1-way ANOVA with Tukey’s post-hoc test. *F* = 4.061. **(L)** No significant differences in levels of p-mTOR (Ser2481) were detected. **(M)** No significant differences in the levels of mTOR were detected.

**Fig. S4.**
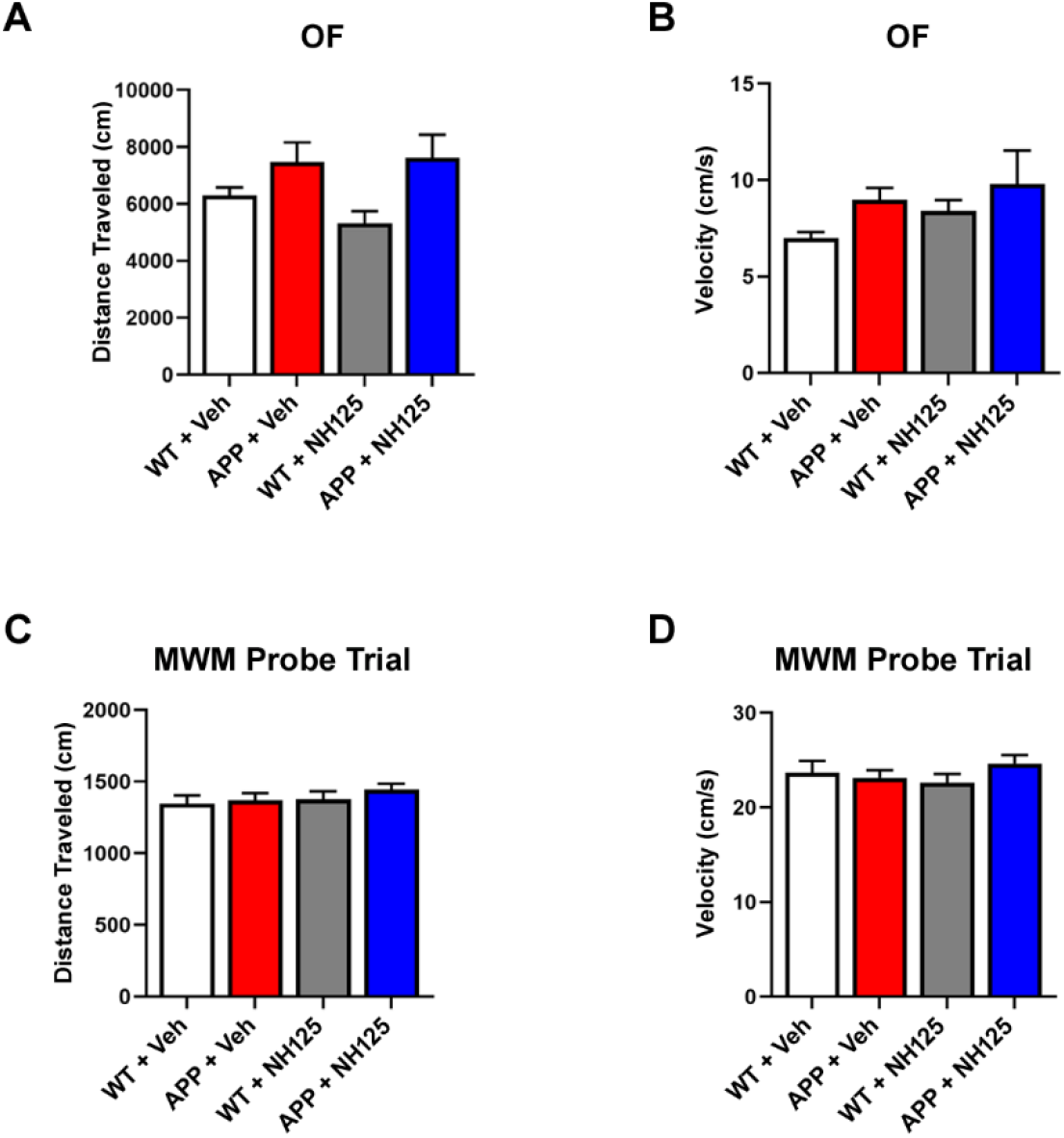
Control behavioral data for APP/PS1 AD model mice treated with NH125. **(A)** Distance traveled in cm of mice in the open field assay (WT + Veh, *n* = 9; APP + Veh, *n* = 10; WT + NH125, *n* = 8; APP + NH125, *n* = 8). **(B)** Velocity of movement in cm/s for open field assay. **(C)** Distance moved in cm for MWM probe trial (WT + Veh, *n* = 13; APP + Veh, *n* = 18; WT + NH125, *n* = 16; APP + NH125, *n* = 16). **(D)** Velocity of movement in cm/s for MWM probe trial.

**Fig. S5.**
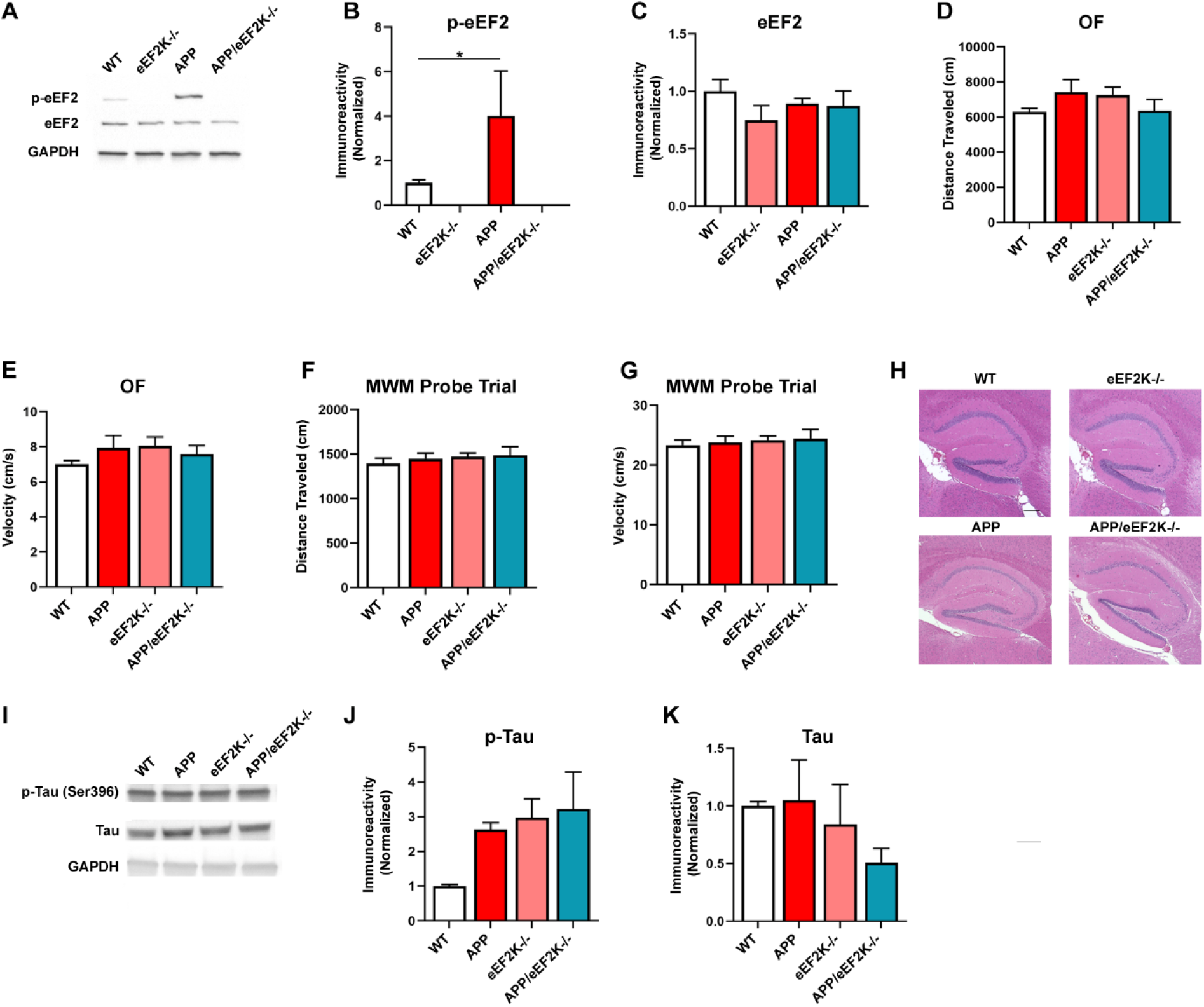
Characterization of phenotypes in APP/PS1 mice with homozygous eEF2K knockout. **(A)** Representative Western blot showing p-eEF2 and eEF2 levels in hippocampal lysates. **(B)** Levels of phosphorylated eEF2 are significantly lower in eEF2K^−/−^ mice than WT mice, and levels of p-eEF2 are significantly lower in APP/eEF2K^−/−^ mice than APP. (WT, *n* = 6; APP, *n* = 5; eEF2K^−/−^, *n* = 4; APP/eEF2K^−/−^ *n* = 6). WT versus eEF2K^−/−^, \**p* < 0.0001; APP versus APP/eEF2K^−/−^, \**p* = 0.03, unpaired *t* tests. **(C)** No differences in total eEF2 levels were detected. **(D)** Distance traveled in cm of mice in the open field assay (WT, *n* = 17; APP, *n* = 9; eEF2K^−/−^, *n* = 10; APP/eEF2K^−/−^, *n* = 8). **(E)** Velocity of movement in cm/s for open field assay. **(F)** Distance traveled in cm for MWM probe trial (WT, *n* = 16; APP, *n* = 9; eEF2K^−/−^, *n* = 16; APP/eEF2K^−/−^, *n* = 9). **(G)** Velocity of movement in cm/s for MWM probe trial. **(H)** Representative images showing hematoxylin & eosin staining in hippocampal sections (20X, scale bar 200 μm). **(I)** Representative Western blots showing levels of phosphorylated tau and tau in hippocampal lysates. **(J)** No significant differences in levels of phosphorylated tau were detected in WT (*n* = 8), APP (*n* = 8), eEF2K^−/−^ (*n* = 6), and APP/eEF2K^−/−^ (*n* = 5) experimental mice. **(K)** No significant differences in levels of total tau were detected.

## Acknowledgements

We thank Dr. George Carlson of McLaughlin Research Institute for providing breeders for Tg19959 AD model mice. We thank Kenneth Grant (Wake Forest School of Medicine) for technical help on TEM imaging. We thank the Wake Forest School of Medicine Pathology Core for their help with tissue processing for immunohistochemistry. We thank the support of the Wake Forest Baptist Comprehensive Cancer Center Proteomics and Metabolomics Shared Resource, supported by the National Cancer Institute’s Cancer Center Support Grant P30CA012197. This work was supported by National Institutes of Health grants R01 AG055581, R01 AG056622 (T.M.), the Alzheimer’s Association grant NIRG-15-362799 (T.M.), the BrightFocus Foundation grant A2017457S (T.M.)

## Author Contributions

Conceptualization: NPK and TM

Methodology: NPK, XZ, XW, WY, HRZ, CEH, EK, HW, AOC, JL, AGR

Investigation: NPK, XZ, XW, WY, HRZ, CEH, EK, HW, AOC, JL, TM

Funding acquisition: CMF and TM

Project administration: NPK, XZ, TM

Supervision: TM

Writing: NPK, AGR, TM

## Competing Interests

The authors declare no competing interests.

## Data and materials availability

All data are available in the main text or the supplementary materials.

## Notes

### Competing Interest Statement

The authors have declared no competing interest.

